# A liver-specific mitochondrial carrier that controls gluconeogenesis and energy expenditure

**DOI:** 10.1101/2022.12.06.519308

**Authors:** Jin-Seon Yook, Zachary H. Taxin, Bo Yuan, Satoshi Oikawa, Christopher Auger, Beste Mutlu, Pere Puigserver, Sheng Hui, Shingo Kajimura

## Abstract

Mitochondria provide essential metabolites and ATP for the regulation of energy homeostasis. For instance, liver mitochondria are a vital source of gluconeogenic precursors under a fasted state. However, the regulatory mechanisms at the level of mitochondrial membrane transport are not fully understood. Here, we report a liver-specific mitochondrial inner-membrane carrier, SLC25A47, which is required for hepatic gluconeogenesis and energy homeostasis. Genome-wide association studies found significant associations between *SLC25A47* and fasting glucose, HbA1c, and cholesterol levels in humans. In mice, we demonstrated that liver-specific deletion of *Slc25a47* impaired hepatic gluconeogenesis selectively from lactate, while significantly enhancing whole-body energy expenditure and the hepatic expression of FGF21. These metabolic changes were not a consequence of general liver dysfunction because acute SLC25A47 deletion in adult mice was sufficient to enhance hepatic FGF21 production, pyruvate tolerance, and insulin tolerance independent of liver damage and mitochondrial dysfunction. Mechanistically, SLC25A47 loss leads to impaired hepatic pyruvate flux and malate accumulation in the mitochondria, thereby restricting hepatic gluconeogenesis. Together, the present study identified a crucial node in the mitochondrial inner-membrane that regulates fasting-induced gluconeogenesis and energy homeostasis.

**SIGNIFICANCE:** Given the impenetrable nature of the mitochondrial inner-membrane, most of the known metabolite carrier proteins, including SLC25A family members, are ubiquitously expressed in mammalian tissues. One exception is SLC25A47 which is selectively expressed in the liver. The present study showed that depletion of SLC25A47 reduced mitochondrial pyruvate flux and hepatic gluconeogenesis under a fasted state, while activating energy expenditure. The present work offers a liver-specific target through which we can restrict hepatic gluconeogenesis, which is often in excess under hyperglycemic and diabetic conditions.

## INTRODUCTION

The role of mitochondria extends far beyond ATP generation. Mitochondria serve as an essential organelle that supplies a variety of important metabolites to the cytosolic compartment and nucleus. An example is in the liver, wherein the mitochondria export phosphoenolpyruvate (PEP) and malate, which serve as gluconeogenic precursors in response to fasting. Under a fed condition, the mitochondria supply citrate that contributes to *de novo* lipogenesis (1, 2). In addition, mitochondria-derived alpha-ketoglutarate (α-KG) functions as a co-factor of Jumonji C domain demethylases (JMJDs) and ten-eleven translocation (TET) enzymes in the nucleus, thereby controlling the transcriptional program, *a.k.a*., retrograde signaling (3).

Mitochondrial flux in the liver is tightly regulated by hormonal cues, such as insulin and glucagon, and dysregulation of these processes profoundly impacts the maintenance of euglycemia, as often seen under the conditions of hyperglycemia and Type 2 diabetes (4–6). For instance, elevated protein expression or activity of pyruvate carboxylase (PC), a mitochondrial matrix-localized enzyme that catalyzes the carboxylation of pyruvate to oxaloacetate (OAA), is associated with hyperglycemia (7, 8). On the other hand, liver-specific deletion of PC potently prevented hyperglycemia in diet-induced obese mice (9). Another example is the mitochondria-localized phosphoenolpyruvate carboxykinase (PCK2, also known as M-PEPCK) that is expressed highly in the liver, pancreas, and kidney, where it catalyzes the conversion of OAA to PEP (10, 11). It has been demonstrated that activation of PCK2 in the liver enhanced the PEP cycle and potentiated gluconeogenesis (12, 13). In turn, depletion of PCK2 in the liver impaired lactate-derived gluconeogenesis and lowered plasma glucose, insulin, and triglycerides in mice (14). Accordingly, a better understanding of mitochondrial metabolite flux in the liver may provide new insights into therapeutic strategies for the management of hyperglycemia and Type 2 diabetes.

Of note, the mitochondrial inner-membrane is impermeable to metabolites relative to the outer membrane. As such, a variety of carrier proteins in the mitochondrial inner-membrane play essential roles in the regulation of metabolite transfer between the matrix and the cytosolic compartment (15, 16). As an example, mitochondrial pyruvate carrier (MPC) mediates the import of pyruvate into the matrix (17, 18). It has been demonstrated that liver-specific deletion of MPC1 or MPC2 reduced mitochondrial TCA flux and impaired pyruvate-driven hepatic gluconeogenesis in diet-induced obese mice (19–21). Recent studies also reported the identification of SLC25A39, which is responsible for glutathione import (22), SLC25A44 for branched-chain amino acids import (23, 24), and SLC25A51 for NAD^+^ import (25–27).

Because of their essential roles, nearly all mitochondrial metabolite carriers (e.g., SLC25A family members) are ubiquitously expressed in mammalian tissues. However, there are two exceptions: Uncoupling protein 1 (UCP1, also known as SLC25A7) that is selectively expressed in brown/beige fat (28), and an orphan carrier, SLC25A47, which is expressed selectively in the liver (see Fig.1A). SLC25A47 was previously described as a mitochondrial protein of which expression was downregulated in hepatocellular carcinoma and that could reduce mitochondrial membrane potential in cultured Hep3B cells, a liver-derived epithelial cell line (29). In yeast, SLC25A47 overexpression elevated mitochondrial electron transport chain uncoupling, implicating its protective role against hepatic steatosis (30). In contrast, a recent study showed that genetic loss of *Slc25a47* led to mitochondrial dysfunction, mitochondrial stress, and liver fibrosis in mice (31). Given these apparently inconsistent reports, this study aims to determine the physiological role of SLC25A47 in systemic energy homeostasis.

**Figure 1.**
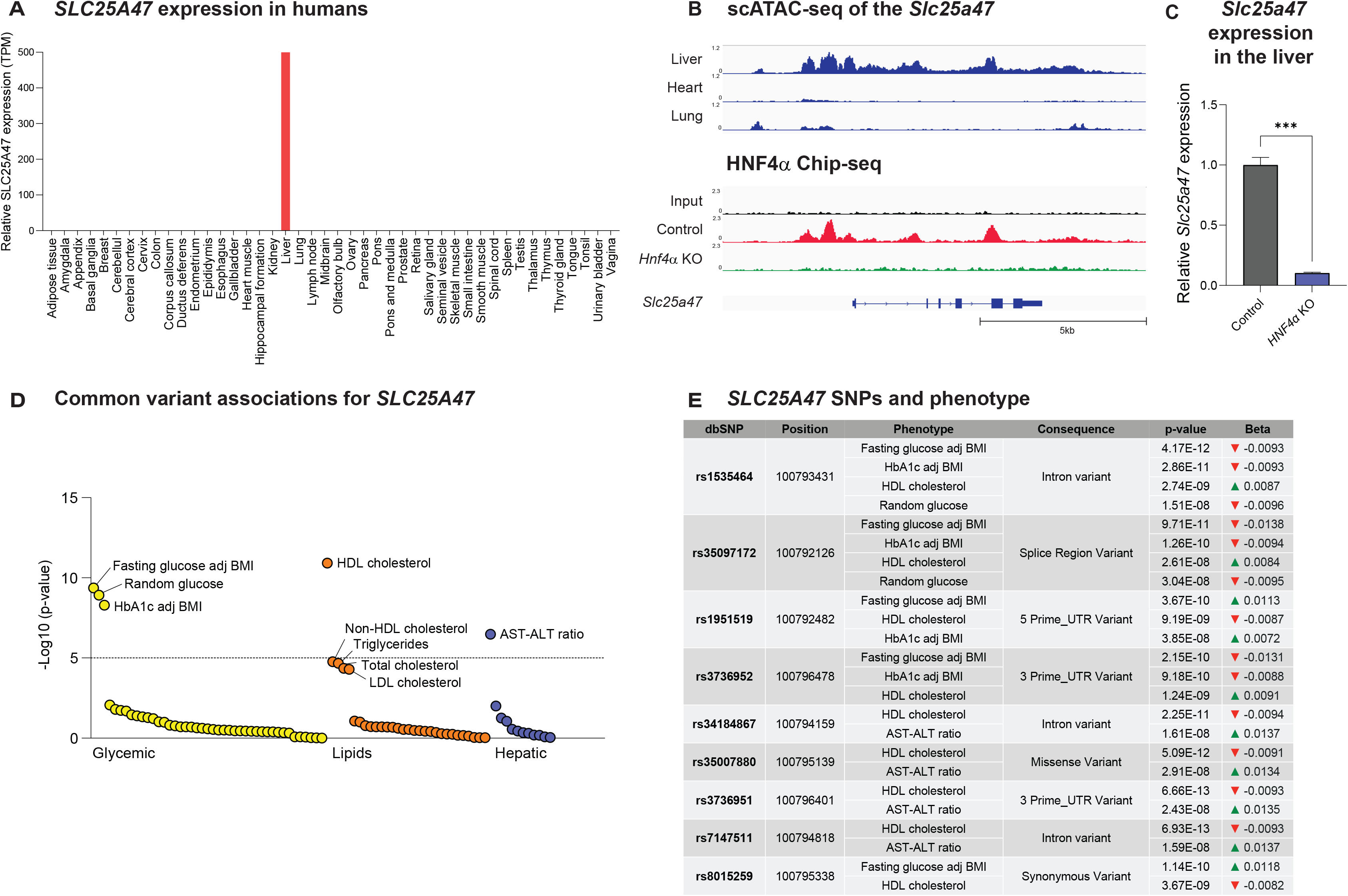
SLC25A47 is a liver-specific mitochondrial carrier that links to human metabolism. **A.** Relative mRNA levels (TPM) in indicated human tissues. The data obtained from Human Protein Atlas (https://www.proteinatlas.org/ENSG00000140107-SLC25A47/tissue) were analyzed. **B. Upper:** The ATAC-seq analysis of the *Slc25a47* gene locus in the liver (upper panel), heart (middle panel), and lung (lower panel). The data were obtained from GEO (GSE111586). **Lower:** The recruitment of HNF4αto the *Slc25a47* gene based on the ChIP-seq data of HNF4α. The data are from ChIP-seq data from GEO (GSE90533). **C.** The mRNA expression of *Slc25a47* in the liver of HNF4 null mouse embryos and controls (18.5-dpc). The data were obtained by analyzing the Affymetrix Mouse Genome 430 2.0 Array data (GSE3126). *n* = 3 for both groups. Data are mean ± SEM.; *** *p* < 0.001 by two-tailed unpaired Student’s *t*-test. **D.** The Phenome-wide association (PheWAS) plot shows significant associations of *SLC25A47* for available traits generated by bottom-line meta-analysis across all datasets in the Common Metabolic Diseases Knowledge Portal (CMDKP). The data can also be acquired from (https://t2d.hugeamp.org/gene.html?gene=SLC25A47) **E.** The genetic associations between *SLC25A47* SNPs and indicated metabolic phenotypes.

## RESULTS

### SLC25A47 is a liver-specific mitochondrial carrier that links to human metabolic disease

The SLC25A solute carrier proteins comprise 53 members in mammals, constituting the largest family of mitochondrial inter-membrane metabolite carriers (32). Among these 53 members, SLC25A47 is unique because this is the sole SLC25A member that is expressed selectively in the liver of mice (29, 31). We independently found that SLC25A47 is selectively expressed in the liver of humans (**Fig. 1A**) and in mice (**Supplementary Fig. S1A**). The publicly available single-cell RNA-seq dataset (33) shows that hepatocytes are the primary cell type that expresses *SLC25A47*, while Kupffer cells also express SLC25A47 that account for approximately 10% of total *Slc25a47* transcripts in the liver (**Supplementary Fig. S1B**).

We next examined the genetic mechanism through which *Slc25a47* is selectively expressed in the liver. The analysis of ATAC-seq data (GSE111586) found an open chromatin architecture in the *Slc25a47* gene locus (chromosome 12: 108,815,740-108,822,741) specific to the liver, whereas the same region appeared to form a heterochromatin structure in the heart and lung (**Fig. 1B, upper**). Notably, the euchromatin region of the *Slc25a47* gene contained binding sites of hepatocyte nuclear factor 4 alpha (HNF4α), to which HNF4αis recruited in the liver (**Fig. 1B, lower**). This result caught our attention because mutations of HNF4αare known to cause MODY1 (maturity-onset diabetes of the young 1), and it plays a central role in the regulation of hepatic and pancreatic transcriptional networks (34, 35). Importantly, HNF4αis required for the hepatic expression of *Slc25a47* as the analysis of a previous microarray dataset (36) found that genetic loss of HNF4αsignificantly attenuated the expression of *Slc25a47* in mouse liver (**Fig. 1C**).

Another important observation is in human genetic association studies from the Type 2 Diabetes Knowledge Portal (type2diabetesgenetics.org), wherein we found significant associations between *SLC25A47* and glycemic and lipid homeostasis. The notable associations include fasting glucose levels adjusted for body mass index (BMI), random glucose levels, HbA1c levels adjusted for BMI, HDL cholesterol levels, and AST-ALT ratio (**Fig. 1D**). One of the strongest single nucleotide polymorphisms (SNIPs) was located in the intronic region of *SLC25A47* (rs1535464) which showed significant associations with lower levels of fasting and random glucose, lower HbA1c levels adjusted for BMI, and higher HDL cholesterol levels (**Fig. 1E**). Similarly, another SNIP (rs35097172) in the regulatory region of *SLC25A47* was associated with lower levels of fasting/random glucose, HbA1c levels adjusted for BMI, and higher HDL cholesterol levels. These data indicate that SLC25A47 is involved in the regulation of glucose and lipid homeostasis, although how these SNPs affect *SLC25A47* expression remains unknown.

### Deletion of *SLC25A47* protects against body-weight gain and lowers plasma cholesterol levels

To determine the physiological role of SLC25A47 in energy homeostasis, we next developed liver-specific SLC25A47 KO mice by crossing *Slc25a47*^flox/flox^ mice with Albumin-Cre (*Alb-Cre*; *Slc25a47*^flox/flox^, herein SLC25A47 KO ^Liver^ mice). We validated that the liver of SLC25A47 KO ^Liver^ mice expressed significantly lower levels of *Slc25A47* mRNA levels than littermate control mice (*Slc25a47*^flox/flox^) by 80 % (**Fig. 2A**). The remaining mRNA in SLC25A47 KO ^Liver^ mice could be attributed to inefficient Cre expression or the transcripts in non-hepatocytes, such as Kupffer cells.

**Figure 2.**
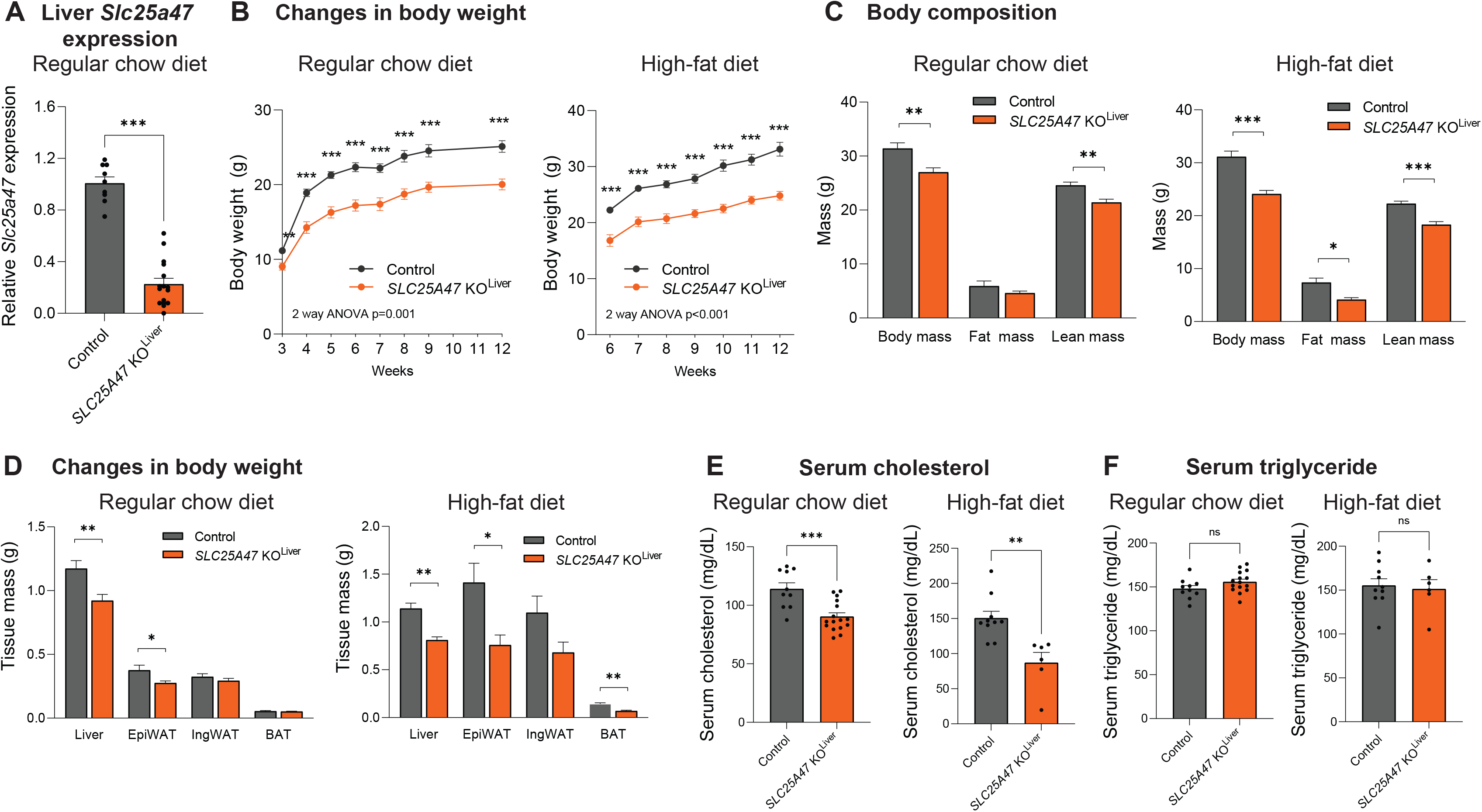
Metabolic characterization of liver-specific SLC25A47 KO mice. **A.** Relative mRNA levels of *Slc25a47* in the liver of *Slc25a47* KO ^Liver^ mice (*Alb-Cre*; *Slc25a47*^lox/flox^) and littermate control mice (*Slc25a47*^lox/flox^). *n* = 10 for *Slc25a47* KO ^Liver^ mice, *n* = 6 for controls, biologically independent mice. Data are mean ± SEM.; *** *p* < 0.001 by two-tailed unpaired Student’s *t*-test. **B.** Changes in body-weight of *Slc25a47* KO ^Liver^ mice and littermate control mice on a regular chow diet (left) and on a high-fat (60%) diet (right). Regular chow diet; *n* = 16 for *Slc25a47* KO ^Liver^ mice on a regular chow diet, *n* = 10 for controls. High fat diet; *n* = 6 for *Slc25a47* KO ^Liver^ mice, *n* = 11 for controls, biologically independent mice. Data are mean ± SEM.; ns, not significant, by two-way repeated-measures ANOVA followed by two-tailed unpaired Student’s *t*-test. **C.** Body composition (fat mass and lean mass) of *Slc25a47* KO ^Liver^ mice and littermate controls at 16 weeks of regular chow diet (left) and at 6 weeks of high-fat diet (right). Regular chow diet; *n* = 11 for *Slc25a47* KO ^Liver^ mice. *n* = 10 for controls. High fat diet; *n* = 6 for *Slc25a47* KO ^Liver^ mice, *n* = 11 for controls, biologically independent mice. Data are mean ± SEM.; ** *p* < 0.01, *** *p* < 0.001 by twotailed unpaired Student’s *t*-test. **D.** Indicated tissue weight of *Slc25a47* KO ^Liver^ mice and littermate control mice in (B). Data are mean ± SEM.; ns, not significant, by two-tailed unpaired Student’s *t*-test. **E.** Serum cholesterol levels of *Slc25a47* KO ^Liver^ mice and littermate control mice at 12 weeks of age on a regular chow diet (left) and after 6 weeks of high-fat diet (right). Regular chow diet; *n* = 16 for *Slc25a47* KO ^Liver^ mice, *n* = 10 for controls. High fat diet; *n* = 6 for *Slc25a47* KO ^Liver^ mice, *n* = 10 for controls. Data are mean ± SEM.; ns, not significant, by two-tailed unpaired Student’s *t*-test. **F.** Serum triglyceride (TG) levels of *Slc25a47* KO ^Liver^ mice and littermate control mice at 12 weeks of age on a regular chow diet in (E). Data are mean ± SEM.; ns, not significant, by two-tailed unpaired Student’s *t*-test.

At birth, there was no difference in the body-weight and body-size between SLC25A47 KO ^Liver^ mice and littermate control mice (**Supplementary Fig. S2A**). However, SLC25A47 KO ^Liver^ mice gained significantly less weight than controls at 3 weeks of age and thereafter on a regular-chow diet (**Fig. 2B, left**). This phenotype was more profound when mice at 6 weeks of age were fed on a high-fat diet (HFD, 60% fat) (**Fig. 2B, right**). The difference in body-weight arose from reduced adipose tissue mass and lean mass both on a regular-chow diet and a high-fat diet (**Fig. 2C**). At tissue levels, adipose tissue and liver mass were lower in SLC25A47 KO ^Liver^ mice relative to control mice (**Fig. 2D**).

Additionally, we found significantly lower serum levels of total cholesterol in SLC25A47 KO ^Liver^ mice than those in controls both on regular-chow and high-fat diets (**Fig. 2E**). On the other hand, we observed no difference in serum triglyceride (TG) levels between the two groups both on regular-chow and high-fat diets (**Fig. 2F**). We found no difference in serum ALT, AST, and albumin levels on a high-fat diet, although serum ALT and AST levels were higher in SLC25A47 KO ^Liver^ mice at 12 weeks of age on a regular-chow diet (**Supplementary Fig. S2B, C, D**).

### Deletion of *SLC25A47* led to elevated whole-body energy expenditure

Given the difference in body-weight between SLC25A47 KO ^Liver^ mice and control mice, we examined the whole-body energy expenditure using metabolic cages. Regression-based analysis of energy expenditure by CaIR-ANCOVA (37) showed that SLC25A47 KO ^Liver^ mice exhibited significantly higher whole-body energy expenditure (kcal/day) independent of body mass at 23°C. The difference remained significant when mice were kept at 30°C (**Fig. 3A**). On the other hand, there was no difference in their food intake and locomotor activity between the genotypes (**Fig. 3B, C**).

**Figure 3.**
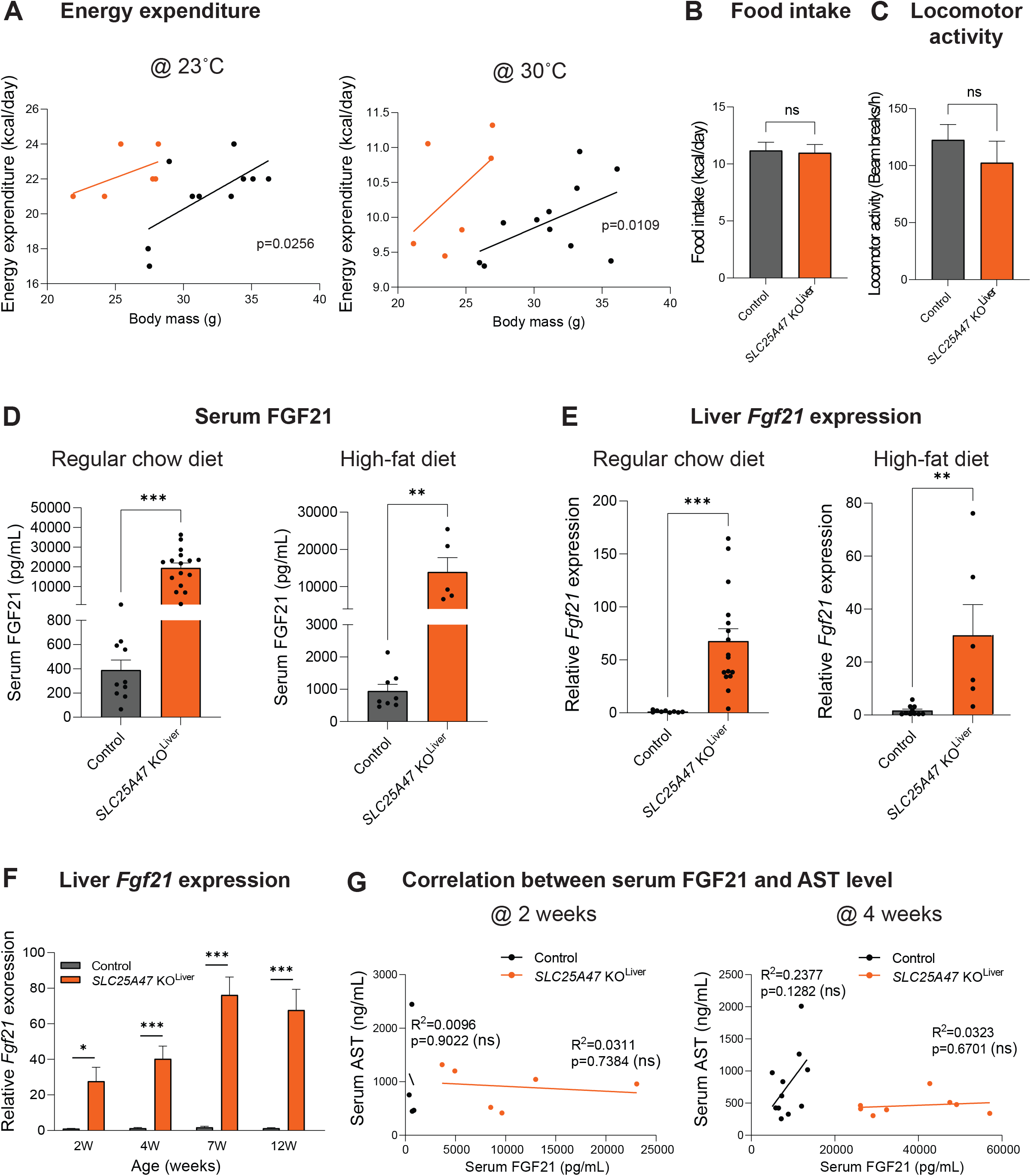
Deletion of *SLC25A47* elevated whole-body energy expenditure. **A.** CaIR-ANCOVA analysis of *Slc25a47* KO ^Liver^ mice and littermate controls at 23°C after one week on a high-fat diet (left) and at 30°C after 3 weeks on a high-fat diet (right). Mice were analyzed by Promethion Metabolic Cage System (Sable Systems). *n* = 6 for *Slc25a47* KO ^Liver^ mice, *n* = 11 for controls. **B.** Food intake of *Slc25a47* KO ^Liver^ mice and littermate control mice at 23°C after one week on a high-fat diet. *n* = 6 for *Slc25a47* KO ^Liver^ mice, *n* = 11 for controls. ns, not significant, by two-tailed unpaired Student’s *t*-test. **C.** Locomotor activity of *Slc25a47* KO ^Liver^ mice and littermate control mice in (B). ns, not significant, by two-tailed unpaired Student’s *t*-test. **D.** Serum levels of FGF21 in *Slc25a47* KO ^Liver^ mice and littermate control mice at 12 weeks of age on a regular chow diet (left) and a high-fat diet for 6 weeks (right). Regular chow diet; *n* = 16 for *Slc25a47* KO ^Liver^ mice, *n* = 10 for controls. High-fat diet; *n* = 16 for *Slc25a47* KO ^Liver^ mice, n = 10 for controls. Data are mean ± SEM.; ns, not significant, by two-tailed unpaired Student’s *t*-test. **E.** Relative expression of *Fgf21* in the liver of *Slc25a47* KO ^Liver^ mice and littermate controls on a regular chow diet (left) and a high-fat diet for 6 weeks (right) in (D). **F.** Relative expression of hepatic *Fgf21* in *Slc25a47* KO ^Liver^ mice and littermate controls on a regular chow diet at indicated ages. *n* = 6 for *Slc25a47* KO ^Liver^ mice, *n* = 4 for controls (2 weeks), *n* = 8 for *Slc25a47* KO ^Liver^ mice, *n* = 14 for controls (4 weeks), *n* = 14 for *Slc25a47* KO ^Liver^ mice, *n* = 14 for controls (7 weeks), *n* = 16 for *Slc25a47* KO ^Liver^ mice, *n* = 9 for controls (12 weeks). Data are mean ± SEM.; ns, not significant, by two-tailed unpaired Student’s *t*-test. **G.** Correlation between serum FGF21 levels and serum AST levels in *Slc25a47* KO ^Liver^ mice and littermate controls at 2 weeks of age (left) and 4 weeks of age (right) on a regular chow diet. *n* = 6 for *Slc25a47* KO ^Liver^ mice, *n* = 4 for controls (2 weeks), *n* = 8 for *Slc25a47* KO ^Liver^ mice, *n* = 11 for controls (4 weeks). Test of the direct association between two variables was performed using simple linear regression.

A possible explanation for the high energy expenditure might be the enhanced thermogenic capacity of brown adipose tissue (BAT) or its sensitivity to β3-adrenergic receptor (β3-AR) signaling. Accordingly, we tested the hypothesis by examining BAT thermogenesis in response to a β3-adrenergic receptor agonist CL316,243 at 30°C. This is a gold-standard method to determine BAT thermogenic responses to β3-AR stimuli, while excluding the contribution of shivering thermogenesis by skeletal muscle (38). We found that a single administration of β3-adrenergic receptor agonist (CL316,243) at 0.5 mg/kg (high dose) potently increased whole-body energy expenditure both in SLC25A47 KO ^Liver^ and littermate controls to a similar degree (**Supplementary Fig. S3A**). This result suggests that the cell-intrinsic thermogenic capacity of BAT, if maximumly activated by a β3-AR stimulus, appears comparable between the two groups. Accordingly, we asked if there was any change in circulating hormonal factors that influenced whole-body energy expenditure of SLC25A47 KO ^Liver^ mice. In this regard, FGF21 is a probable candidate because it is a well-established endocrine hormone that increases energy expenditure by activating the sympathetic nervous system (39). Consistent with the recent work (31), we found that serum levels of FGF21 in SLC25A47 KO ^Liver^ mice were significantly higher relative to littermate controls both on regular-chow and high-fat diets (**Fig. 3D**). The increase in circulating FGF21 levels was due to elevated *Fgf21* transcription in the liver (**Fig. 3E**). This is in agreement with the previous work demonstrating that the liver is the primary source of circulating FGF21 (40).

Of note, elevated *Fgf21* gene expression in SLC25A47 KO ^Liver^ mice was already observed at 2 weeks of age, a time point in which there was no difference in body-weight, serum ATL/AST levels, and mitochondrial stress-related genes in the liver (**Fig. 3F, Supplementary Fig. S3B, C, D**). Importantly, there was no correlation between serum FGF21 levels and AST levels in control and SLC25A47 KO ^Liver^ mice at 2 and 4 weeks of age (**Fig. 3G**). The results indicate that the stimulatory effect of SLC25A47 deletion on FGF21 expression is not merely a consequence of liver damage. We addressed this point further in the following sections (see Fig. 5).

### SLC25A47 is required for pyruvate-derived hepatic gluconeogenesis *in vivo*

We next examined the extent to which SLC25A47 regulates systemic glucose homeostasis. This is based on the observation that fasting glucose levels of SLC25A47 KO ^Liver^ mice were consistently lower than littermate controls both on regular-chow and high-fat diets (**Fig. 4A**). At 4 weeks of high-fat diet, we found no major difference in glucose tolerance between the two groups, although fasting glucose levels were lower in SLC25A47 KO ^Liver^ mice than control mice (**Fig. 4B**). In contrast, SLC25A47 KO ^Liver^ mice exhibited significantly higher insulin tolerance than controls in response to insulin at a low dose (0.4 U/kg) (**Fig. 4C**). It is notable that SLC25A47 KO ^Liver^ mice remained hypoglycemic (< 70 mg/dl) following insulin administration.

**Figure 4.**
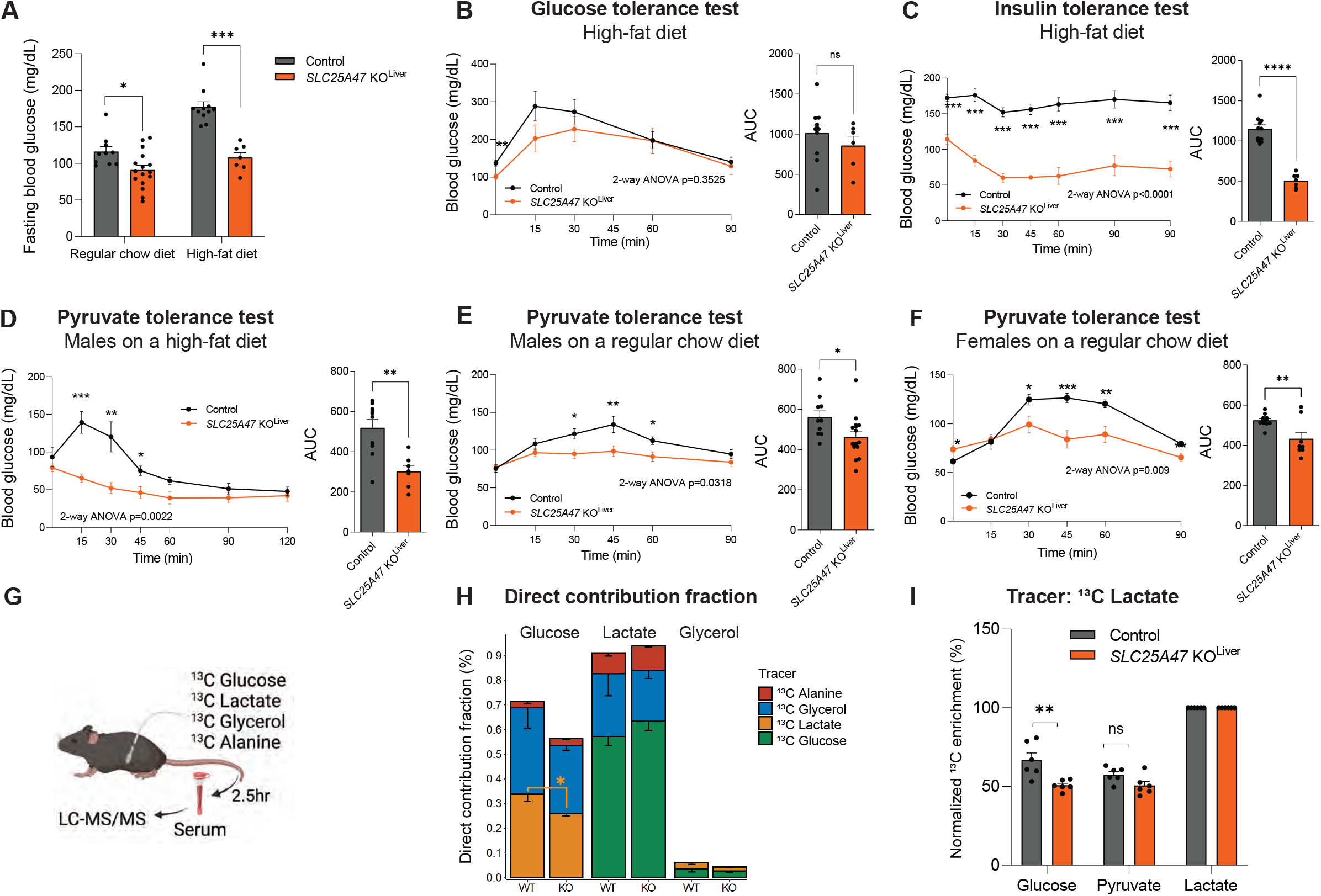
SLC25A47 is required for pyruvate-derived hepatic gluconeogenesis. **A.** Fasting blood glucose levels (6 hours) in *Slc25a47* KO ^Liver^ mice and littermate controls at 7 weeks of age on a regular chow diet (left) and at 4 weeks on a high-fat diet (right). Regular diet; *n* = 16 for *Slc25a47* KO ^Liver^ mice, *n* = 10 for control mice. High fat diet; *n* = 6 for *Slc25a47* KO ^Liver^ mice, *n* = 11 for control mice. **B.** Glucose tolerance test in *Slc25a47* KO ^Liver^ mice and littermate controls at 4 weeks of high-fat diet. After 6 hours of fasting, mice received *i.p*. injection of glucose at 2 g kg^-1^ body-weight. *n* = 6 for *Slc25a47* KO ^Liver^ mice, *n* = 11 for control mice, biologically independent mice. Data are mean ± SEM.; *P*-value was determined by two-way repeated-measures ANOVA followed by Fisher’s LSD test. Right: Area under the curve (AUC) of the data was calculated by Graphpad software. ns, not significant by unpaired Student’s *t*-test. **C.** Insulin tolerance test in *Slc25a47* KO ^Liver^ mice and littermate controls at 4 weeks of high-fat diet. After 6 hours of fasting, mice received *i.p*. injection of insulin at 0.4 U kg^-1^ body-weight. *n* = 7 for *Slc25a47* KO ^Liver^ mice, *n* = 11 for control mice, biologically independent mice. Data are mean ± SEM.; *P*-value was determined by two-way repeated-measures ANOVA followed by Fisher’s LSD test. Right: AUC of the data. **** *p* < 0.0001 by two-tailed unpaired Student’s *t*-test. **D.** Pyruvate tolerance test in *Slc25a47* KO ^Liver^ mice and littermate controls on a high-fat diet for 3 weeks. After 16 hours of fasting, mice received *i.p*. injection of pyruvate at 2 g kg^-1^ body-weight. *n* = 6 for *Slc25a47* KO ^Liver^ mice, *n* = 11 for control mice, biologically independent mice. Data are mean ± SEM.; *P*-value was determined by two-way repeated-measures ANOVA followed by Fisher’s LSD test. Right: AUC of the data was calculated by Graphpad software. ** *p* < 0.01 by two-tailed unpaired Student’s *t*-test. **E.** Pyruvate tolerance test in male *Slc25a47* KO ^Liver^ mice and littermate controls on a regular chow diet. After 16 hours of fasting, mice received *i.p*. injection of pyruvate at 2 g kg^-1^ body-weight. *n* = 16 for *Slc25a47* KO ^Liver^ mice, *n* = 10 for control mice, biologically independent mice. AUC of the data was calculated by Graphpad software. * *p* < 0.05 by two-tailed unpaired Student’s *t*-test. **F.** Pyruvate tolerance test in female *Slc25a47* KO ^Liver^ mice and littermate controls on a regular chow diet. After 16 hours of fasting, mice received *i.p*. injection of pyruvate at 2 g kg^-1^ body-weight. *n* = 8 for *Slc25a47* KO ^Liver^ mice, *n* = 12 for control mice, biologically independent mice. AUC of the data was calculated by Graphpad software. ** *p* < 0.01 by two-tailed unpaired Student’s *t*-test. **G.** Schematic illustration of tracer experiments. *Slc25a47* KO ^Liver^ mice and littermate controls on a regular chow diet were infused with indicated ^13^C-labeled tracer via the catheter. During the infusion, we collected serum from fasted mice and analyzed by LC-MS/MS. **H.** The relative contribution of ^13^C-labeled tracers to glucose, lactate, and glycerol in (G). *n* = 6 for *Slc25a47* KO ^Liver^ mice, *n* = 6 for control mice, biologically independent mice. Data are mean ± SEM.; * *p* < 0.05 by two-tailed unpaired Student’s *t*-test. **I.** Direct contribution of ^13^C-labeled lactate to circulating levels of glucose, pyruvate, and lactate in (G).

Pyruvate tolerance tests found that SLC25A47 KO ^Liver^ mice at 3 weeks of high-fat diet exhibited significantly lower hepatic gluconeogenesis than control mice (**Fig. 4D**). Of note, the difference in pyruvate tolerance was independent of diet and sex, as we observed consistent results both in male and female mice on a regular-chow diet (**Fig. 4E, F**). On the other hand, there was no difference in glucose-stimulated serum insulin levels and hepatic glycogen contents between the two groups (**Supplementary Fig. S4A, B**). These results led to the hypothesis that the lower fasting glucose levels seen in SLC25A47 KO ^Liver^ mice are attributed to reduced hepatic gluconeogenesis, rather than impaired glycogenolysis or elevated insulin sensitivity in the skeletal muscle.

To test the hypothesis, we next examined the contribution of hepatic gluconeogenesis to circulating glucose by infusing fasted mice with U-^13^C-labeled lactate or ^13^C-labeled glucose. To examine the relative contribution of other gluconeogenic precursors to blood glucose, we also infused fasted mice with U-^13^C-labeled glycerol and U-^13^C-alanine (**Fig. 4G**). During the infusion, we collected and analyzed serum from fasted mice using liquid-chromatography-mass spectrometry (LC-MS), as described in recent studies (41, 42). We used ^13^C-lactate as a gluconeogenic precursor instead of pyruvate because circulating lactate is the primary contributor to gluconeogenesis and in rapid exchange with pyruvate (43).

The analyses showed that glucose production from ^13^C-lactate was significantly lower in SLC25A47 KO ^Liver^ mice than in control mice (**Fig. 4H** orange bars). Notably, this impairment was selective to the lactate-to-glucose conversion, as we found no significant difference in glucose production from ^13^C-glycerol between the two groups (**Fig. 4H** blue bars, **Supplementary Fig. S4C**). The relative contribution of alanine to serum glucose was far less than lactate, with no statistical difference between the genotypes (**Fig. 4H** red bars). The lactate-to-pyruvate conversion was unaffected in SLC25A47 KO ^Liver^ mice, suggesting that impaired gluconeogenesis from lactate is attributed to reduced pyruvate utilization in the liver (**Fig. 4I**). We also found no difference in the conversion from ^13^C-glucose to pyruvate and lactate (**Supplementary Fig. S4D**). These results indicate that SLC25A47 is required selectively for gluconeogenesis from lactate under a fasted condition, whereas it is dispensable for gluconeogenesis from other substrates.

### Acute depletion of SLC25A47 improved glucose homeostasis without causing liver damage

Recent work by Brescinani et al. suggested the possibility that the metabolic changes in SLC25A47 KO ^Liver^ mice, such as elevated FGF21 and impaired glucose production, were merely secondary to general hepatic dysfunction and fibrosis (31). To exclude metabolic complications caused by chronic deletion of SLC25A47, particularly during the prenatal and early postnatal periods, we aimed to acutely deplete SLC25A47 in adult mice. To this end, we acutely depleted SLC25A47 in adult mice by delivering AAV-TGB-Cre or AAV-TGB-null (control) into the liver of *Slc25a47*^flox/flox^ mice via tail-vein (**Fig.5A**). AAV-Cre administration successfully reduced *Slc25a47* mRNA expression by approximately 50% (**Fig. 5B**). Although the depletion efficacy of AAV-Cre was less than the genetic approach using *Albumin*-Cre, this model gave us an opportunity to determine the extent to which acute and partial depletion of SLC25A47 in adult mice sufficiently affect hepatic glucose production and energy expenditure, while avoiding metabolic complications associated with chronic SLC25A47 deletion.

**Figure 5.**
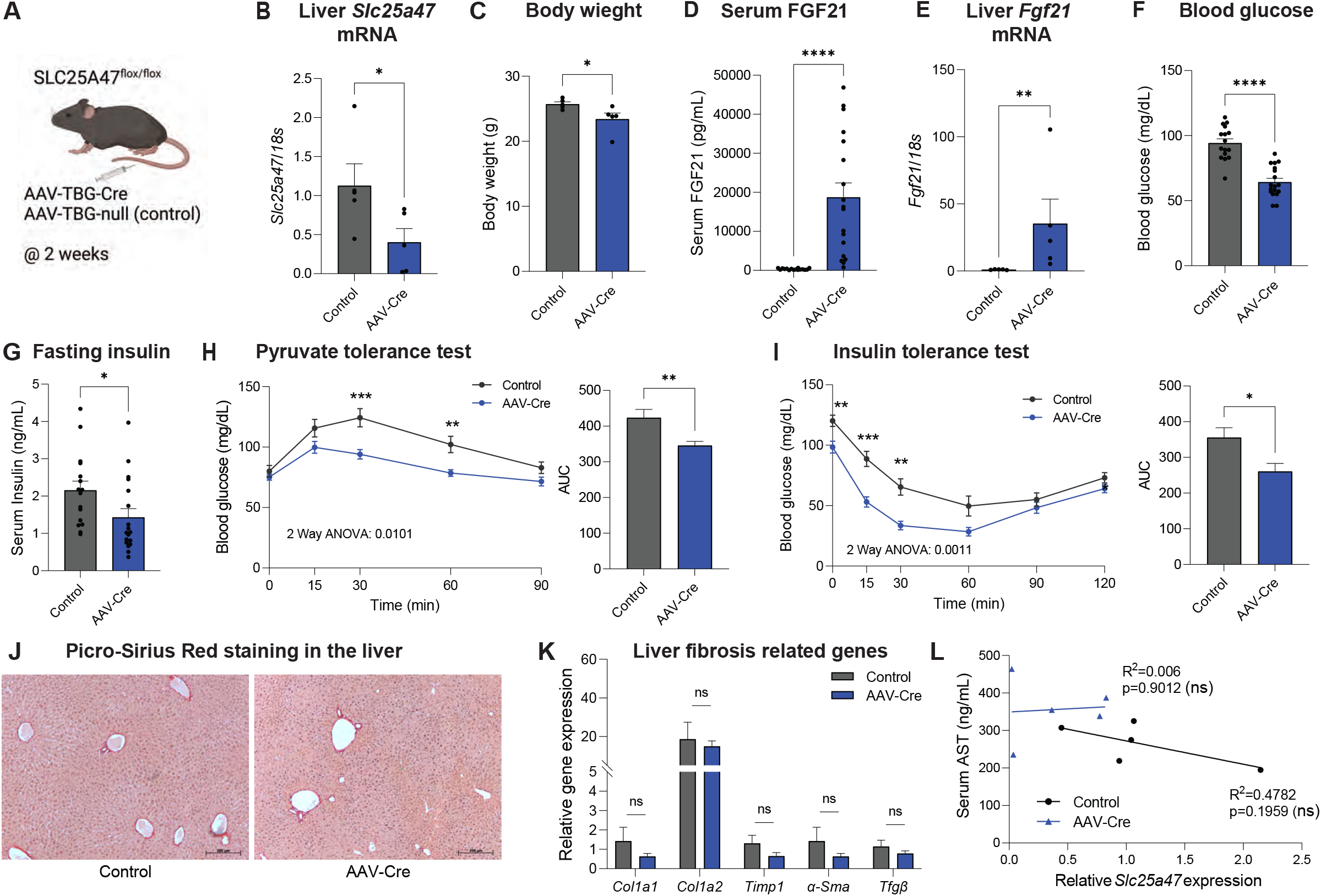
Acute depletion of SLC25A47 improves glucose homeostasis independent of liver damage. **A.** Schematic illustration of acute depletion of *Slc25a47* study. *Slc25a47^flox/flox^* mice at 7 weeks of age on a regular chow diet received AAV-Cre or AAV-null (control) via tail-vein. **B.** Relative mRNA levels of hepatic *Slc25a47* in *Slc25a47*^flox/flox^ mice that received AAV-Cre or AAV-null (control) via tail vein. Samples were collected at 2 weeks after AAV injection. *n* = 5 for controls, *n* = 5 for AAV-Cre, biologically independent mice. Data are mean ± SEM.; * *p* < 0.05 by Mann-Whitney test. **C.** Body weight of *Slc25a47*^flox/flox^ mice at 2 weeks after receiving AAV-Cre or AAV-null (control). Mice were on a regular chow diet. *n* = 5 for controls, *n* = 5 for AAV-Cre, biologically independent mice. * *p* < 0.05 by two-tailed unpaired Student’s *t*-test. **D.** Serum FGF21 levels in *Slc25a47*^flox/flox^ mice at 2 weeks after receiving AAV-Cre or AAV-null (control) Mice were on a regular chow diet. *n* = 16 for controls, *n* = 18 for AAV-Cre, biologically independent mice. **** *p* < 0.0001 by two-tailed unpaired Student’s *t*-test. **E.** Relative expression of hepatic *Fgf21* in *Slc25a47*^flox/flox^ mice at 2 weeks after receiving AAV-Cre or AAV-null (control). *n* = 5 for controls, *n* = 5 for AAV-Cre, biologically independent mice. ** *p* < 0.01 by Mann-Whitney test. **F.** Fasting blood glucose levels (6 hours) in *Slc25a47*^flox/flox^ mice at 2 weeks after receiving AAV-Cre or AAV-null (control). *n* = 5 for controls, *n* = 5 for AAV-Cre, biologically independent mice. **** *p* < 0.0001 by two-tailed unpaired Student’s *t*-test. **G.** Fasting insulin levels in *Slc25a47*^flox/flox^ mice at 2 weeks after receiving AAV-Cre or AAV-null (control). *n* = 16 for controls, *n* = 18 for AAV-Cre, biologically independent mice. * *p* < 0.05 by twotailed unpaired Student’s *t*-test. **H.** Pyruvate tolerance test in *Slc25a47*^flox/flox^ mice at 2 weeks after receiving AAV-Cre or AAV-null (control). After 16 hours of fasting, mice received *i.p*. injection of pyruvate at 2 g kg^-1^ body-weight. *n* = 15 for controls, *n* = 16 for AAV-Cre, biologically independent mice. Data are mean ± SEM.; *P*-value was determined by two-way repeated-measures ANOVA followed by Fisher’s LSD test. Right: Area under the curve (AUC) of the data was calculated by Graphpad software. ** *p* < 0.01 by twotailed unpaired Student’s *t*-test. **I.** Insulin tolerance test in *Slc25a47*^flox/flox^ mice at 6 weeks after receiving AAV-Cre or AAV-null (control). After 6 hours of fasting, mice received *i.p*. injection of insulin at 0.4 U kg^-1^ body-weight. *n* = 11 for controls, *n* = 13 for AAV-Cre, biologically independent mice. Right: AUC of the data. * *p* < 0.05 by two-tailed unpaired Student’s *t*-test. **J.** Representative image of Picro-Sirius Red staining in the liver of *Slc25a47*^flox/flox^ mice at 2 weeks after receiving AAV-Cre or AAV-null (control). Scale = 200 μm. **K.** Relative mRNA levels of indicated fibrosis marker genes in the liver of *Slc25a47*^flox/flox^ mice that received AAV-Cre or AAV-null (control). Samples were collected at 2 weeks after AAV injection. *n* = 5 for controls, *n* = 5 for AAV-Cre, biologically independent mice. Data are mean ± SEM.; ns, not significant, by two-tailed unpaired Student’s *t*-test. **L.** Correlation between serum AST levels and hepatic *Slc25a47* expression in *Slc25a47*^flox/flox^ mice at 2 weeks after receiving AAV-Cre or AAV-null (control). *n* = 5 for controls, *n* = 5 for AAV-Cre, biologically independent mice. ns, not significant

After 2 weeks of AAV administration, we found that acute SLC25A47 depletion led to reduced body-weight gain (**Fig. 5C, Supplementary Fig. S5A**) and increased serum FGF21 levels (**Fig. 5D**). The increase in serum FGF21 levels was associated with elevated hepatic FGF21 mRNA expression (**Fig. 5E**). Consistent with the observations in liver-SLC25A47 KO mice, acute SLC25A47 depletion resulted in reduced fasting serum glucose levels (**Fig. 5F**) and insulin levels (**Fig. 5G**). Importantly, acute SLC25A47 depletion improved systemic pyruvate tolerance (**Fig. 5H**) and insulin tolerance (**Fig. 5I**). In contrast, acute SLC25A47 did not alter systemic glycerol tolerance, although there was a modest change at later time points after glycerol administration (**Supplementary Fig. S5B**). The difference in glycerol tolerance at later time points is likely because glycerol-derived glucose is converted to lactate in peripheral tissues, which is eventually utilized as a gluconeogenic substrate (44).

Next, we examined if such metabolic changes were associated with liver injury *in vivo*. Histological analyses by Picro-Sirius Red staining did not find any noticeable sign of liver fibrosis (**Fig. 5J**). Similarly, histological analyses by hematoxylin and eosin (H&E) staining found no difference between control vs. AAV-Cre injected mice (**Supplementary Fig. S5C**). Furthermore, acute SLC25A47 depletion did not alter the expression of liver fibrosis marker genes (**Fig. 5K**). Also, we found no significant correlation between serum AST levels and hepatic SLC25A47 expression (**Fig. 5L**) and between serum AST levels and FGF21 levels (**Supplementary Fig. S5D**). Moreover, we observed no significant difference in the Complex I and II activities of isolated liver mitochondria between the two groups (**Supplementary Fig. S5E**). These data suggest that acute SLC25A47 depletion sufficiently enhanced hepatic FGF21 expression, pyruvate tolerance, and insulin tolerance independent of liver damage and hepatic mitochondrial dysfunction.

### SLC25A47 is required for mitochondrial pyruvate flux and malate export

We next asked which steps of the lactate-derived hepatic gluconeogenesis were altered in SLC25A47 KO ^Liver^ mice. To this end, we took unbiased omics approaches – RNA-seq and mitochondrial metabolomics analyses – in the liver of SLC25A47 KO ^Liver^ mice and littermate controls under a fasted condition. The summary of the results is shown in **Fig. 6A**. The RNA-seq data analysis found that the liver of SLC25A47 KO ^Liver^ mice expressed significantly higher levels of *Pkm, Eno3, Aldoa, Fbp1, Gpi1*, and *G6pc3* (**Fig. 6B**), suggesting a compensatory upregulation of gluconeogenic gene expression in SLC25A47 KO ^Liver^ mice.

**Figure 6.**
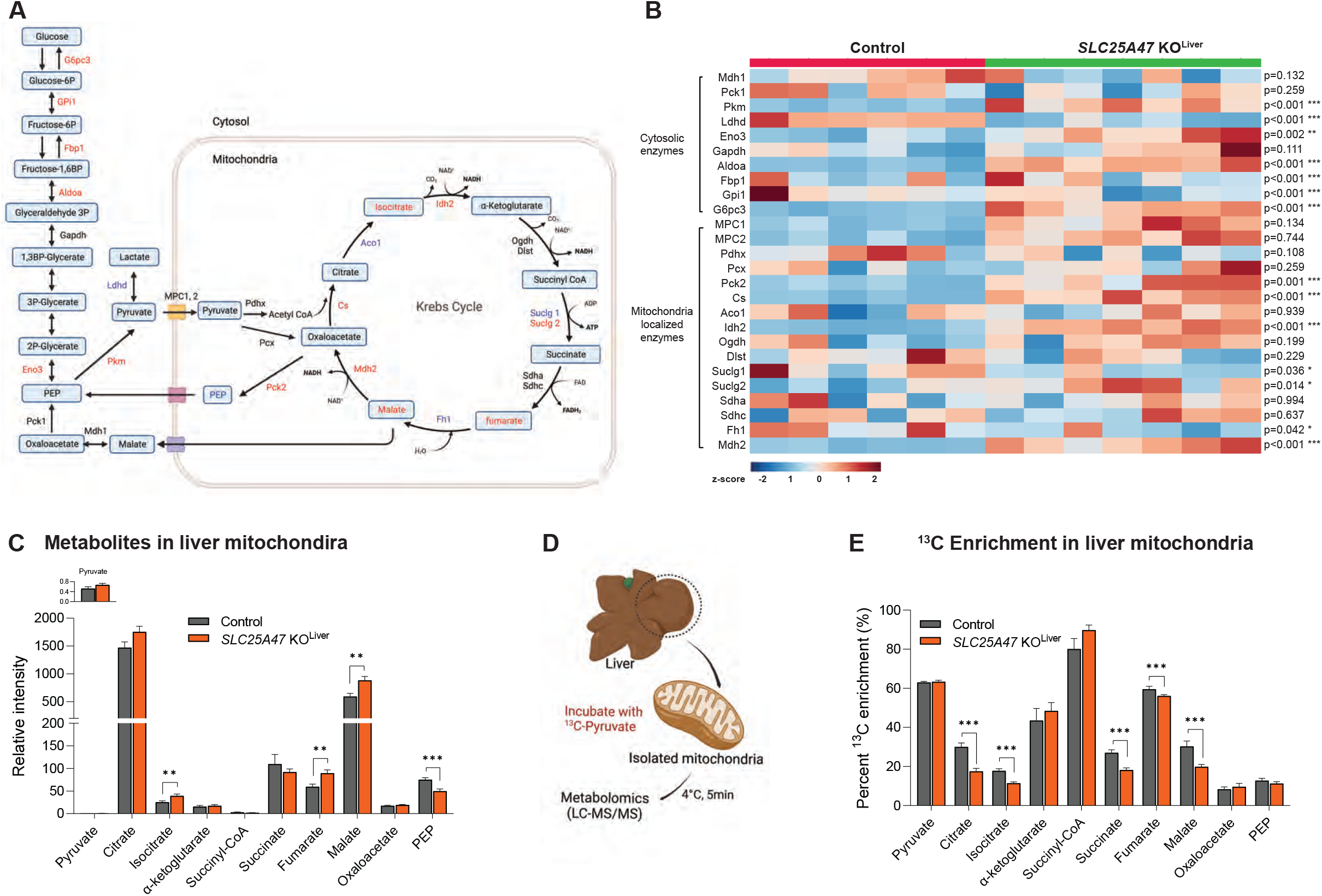
SLC25A47 is required for mitochondrial pyruvate efflux and malate export. **A.** Summary of RNA-seq and mitochondrial metabolomics data in the liver of *Slc25a47* KO ^Liver^ mice and littermate controls at 12 weeks of age on a regular chow diet. Red letters indicate upregulated in *Slc25a47* KO ^Liver^ mice relative to controls. Blue letters indicate downregulated in *Slc25a47* KO ^Liver^ mice. Black letters indicate no statistical changes between the genotypes. **B.** Heatmap of mRNA levels of indicated cytosolic gluconeogenic enzymes and mitochondria localized enzymes in the liver of *Slc25a47* KO ^Liver^ mice and control mice on a regular chow diet. Mice at 12 weeks of age have fasted for 6 hours. *n* = 7 for *Slc25a47* KO ^Liver^ mice, *n* = 6 for control mice, biologically independent mice. The color scale shows *Z*-scored Transcripts Per Million-values (TPM) representing the mRNA level of each gene in the blue (low expression)-red (high expression) scheme. *p*-value was determined by two-tailed unpaired Student’s *t*-test. **C.** Relative levels of indicated metabolites in the liver mitochondria of *Slc25a47* KO ^Liver^ mice and control mice on a regular chow diet. Mice at 7 weeks of age fasted for 6 hours. *n* = 14 for *Slc25a47* KO ^Liver^ mice, *n* = 14 for control mice, biologically independent mice. Data were normalized to mitochondrial protein levels (ug/mL) and shown as mean ± SEM.; *P*-value was determined by twotailed unpaired Student’s *t*-test. **D.** Schematic illustration of tracer experiments in isolated mitochondria. Mitochondria were isolated from the liver of fasted mice. Isolated mitochondria were incubated with ^13^C-labeled pyruvate. Subsequently, mitochondrial metabolites were analyzed by LC-MS/MS. **E.** Direct contribution of ^13^C-labeled pyruvate to indicated metabolites in the mitochondria. *n* = 14 for *Slc25a47* KO ^Liver^ mice, *n* = 14 for littermate control mice, biologically independent mice. Data were normalized to mitochondrial protein (ug/mL) and shown as mean ± SEM., ns, not significant, by twotailed unpaired Student’s *t*-test.

A notable finding is the distinct regulation of mitochondrial matrix-localized enzymes vs. cytosolic enzymes: we found that the expression of the mitochondrial TCA cycle enzymes, such as citrate synthase (*Cs*), the mitochondrial form of isocitrate dehydrogenase (*Idh2*), and *Suclg2* (the subunits of succinate-CoA ligase) was significantly upregulated in the liver of SLC25A47 KO ^Liver^ mice relative to controls. In addition, the expression of *Pck2*, the mitochondria-localized PEPCK that converts OAA to PEP within the mitochondria, was upregulated in the liver of SLC25A47 KO ^Liver^ mice. In contrast, the expression of the cytosolic form of PEPCK (*Pck1*) was unchanged. Similarly, the expression of *Mdh2*, which catalyzes the conversion between oxaloacetate and malate in the mitochondria, was significantly elevated in the liver of SLC25A47 KO ^Liver^ mice, whereas the expression of *Mdh1*, the cytosolic form, showed a trend of down-regulation. These results suggest that SLC25A47 loss leads to a distinct gene expression pattern of mitochondrial vs. cytosolic enzymes that control hepatic gluconeogenesis.

The mitochondrial metabolomics analysis revealed that the liver mitochondria of SLC25A47 KO ^Liver^ mice accumulated significantly higher levels of isocitrate, fumarate, and malate than that of control mice (**Fig. 6C**). In contrast, mitochondrial PEP contents were lower in SLC25A47 KO livers relative to controls. We found no difference in the mitochondrial contents of pyruvate, citrate, α-KG, succinyl CoA, succinate, and OAA between the two groups. Additionally, there was no difference in the mitochondrial contents of co-factors required for the TCA cycle reactions, such as Coenzyme A, NADH, NADP^+^, NADPH, and FAD, although mitochondrial NAD^+^ and GTP levels were higher in SLC25A47 KO ^Liver^ mice than controls (**Supplementary Fig. 6A**).

The above data led to the hypothesis that SLC25A47 controls either pyruvate import to the mitochondrial matrix or pyruvate flux within the mitochondria. To test this, we isolated mitochondria from the liver of SLC25A47 KO ^Liver^ mice and littermate controls under a fasted condition. The isolated mitochondria were incubated with [U-^13^C] labeled pyruvate and subsequently analyzed by LC-MS/MS (**Fig. 6D**). We found no difference in the mitochondrial contents of ^13^C-pyruvate levels between the two groups, suggesting that mitochondrial pyruvate uptake *per se* was not altered in the liver of SLC25A47 KO ^Liver^ mice (**Fig. 6E**). This is in agreement with the data that the expression of MPC1 and MPC2 was not different between the genotypes (see Fig. 6B). On the other hand, the enrichments of ^13^C-labeled citrate, isocitrate, succinate, fumarate, and malate were significantly lower in the mitochondria of SLC25A47 KO ^liver^ mice than those in controls (**Fig. 6E**). There was no difference in ^13^C-labeled OAA and PEP between the groups. Together, these results suggest that genetic loss of SLC25A47 impaired mitochondrial pyruvate flux, leading to an accumulation of fumarate, malate, and isocitrate in the liver mitochondria. Impaired export of malate from the mitochondria into the cytosolic compartment leads to reduced lactate-derived hepatic gluconeogenesis under a fasted condition.

## DISCUSSION

Mitochondrial flux in the liver is highly nutrition-dependent. Under a fed condition, malate is imported into the mitochondrial matrix in exchange for α-KG *via* mitochondrial α-KG/malate carrier (SLC25A11) as a part of the malate-aspartate shuttle, a mechanism to transport reducing equivalents (NADH) into the mitochondrial matrix (45). In addition, mitochondrial dicarboxylate carrier SLC25A10 can mediate the import of malate into the mitochondrial matrix in addition to malonate, succinate, phosphate, sulfate, and thiosulfate (46). Under a fasted state, when liver glycogen is depleted, malate is exported from the mitochondrial matrix into the cytosolic compartment, where it is converted to OAA by MDH1 and utilized as a gluconeogenic substrate. However, what controls the nutrition-dependent mitochondrial malate flux remains elusive. The present work showed that SLC25A47 loss led to an accumulation of mitochondrial malate and reduced hepatic gluconeogenesis, without affecting gluconeogenesis from glycerol. The results indicate that SLC25A47 mediates the export of mitochondria-derived malate into the cytosol. However, the present study could not exclude the possibility that SLC25A47 mediates the transport of co-factors needed for mitochondrial pyruvate flux, although we found no difference in the mitochondrial contents of Coenzyme A and NADH between the genotypes. Our future study aims to determine the specific substrate of SLC25A47 by biochemically reconstituting this protein in a cell-free system, such as liposomes.

The present work showed that genetic loss of SLC25A47 reduced mitochondrial pyruvate flux, thereby restricting lactate-derived hepatic gluconeogenesis and preventing hyperglycemia. This is in alignment with several mouse models with impaired mitochondrial pyruvate flux in the liver. For instance, liver-specific loss of pyruvate carboxylase (PC) limits the supply of pyruvate-derived OAA in the mitochondria, leading to reduced TCA flux and hepatic gluconeogenesis (9). Similarly, liver-specific deletion of the mitochondrial pyruvate carrier (MPC1 or MPC2) or the mitochondria-localized PEPCK (M-PEPCK) reduces hepatic gluconeogenesis and protects mice against diet-induced hyperglycemia (14, 19–21). A recently developed non-invasive method, Positional Isotopomer NMR tracer analysis (PINTA), would be instrumental to determine how SLC25A47 loss alters the rates of hepatic mitochondrial citrate synthase flux vs. pyruvate carboxylase flux (35).

It is worth pointing out that elevated energy expenditure and reduced body-weight are unique to SLC25A47 KO mice. Indeed, no changes in energy expenditure and body-weight were seen in liver-specific MPC1/2 KO mice or M-PEPCK KO mice relative to the respective control mice. Elevated energy expenditure of SLC25A47 KO mice appears to be attributed to elevated FGF21 because recent work demonstrated that deletion of FGF21 abrogated the effects of SLC25A47 on energy expenditure and body-weight (31). Importantly, our results suggest that partial SLC25A47 depletion was sufficient to stimulate FGF21 production independently from liver damage. It is conceivable that changes in mitochondria-derived metabolites, such as malate and others, control the transcription of FGF21 *via* retrograde signaling (3). Our future study will explore the mechanisms through which SLC25A47-mediated mitochondrial signals control the nuclear-coded transcriptional program in a nutrition-dependent manner.

With these results in mind, we consider that SLC25A47 is a plausible target for hyperglycemia and Type 2 diabetes for the following reasons. First, excess hepatic gluconeogenesis is commonly seen in human hyperglycemia and Type 2 diabetes (4–6). Notably, GWAS data found significant associations between *SLC25A47* and glycemic homeostasis in humans – particularly, several SNPs in the *SLC25A47* were significantly associated with lower levels of glucose and HbA1c adjusted for BMI, although how these SNPs affect *SLC25A47* expression awaits future studies. Second, SLC25A47 is exceptionally unique among 53 members of the mitochondrial SLC25A carriers, given its selective expression in the liver. This tissue-specificity makes SLC25A47 an attractive therapeutic target, considering the recent successful examples in which liver-targeting mitochondrial uncouplers protected mice against Type 1 and Type 2 diabetes, hepatic steatosis, and cardiovascular complications (47–49). A potential caveat is the detrimental effect associated with chronic SLC25A47 deletion, such as mitochondrial stress, lipid accumulation, and fibrosis (31). However, our data showed that acute depletion of SLC25A47 by ~50% sufficiently restricted gluconeogenesis and enhanced insulin tolerance in adult mice without causing liver fibrosis and mitochondrial dysfunction. Thus, it is conceivable that temporal and partial inhibition of SLC25A47 by using small-molecule inhibitors or anti-sense oligos would be effective in restricting excess hepatic gluconeogenesis while avoiding the detrimental side effects.

## ACKNOWLEDGEMENT

We are grateful to Drs. Jose C. Florez and Maria Costanzo at Massachusetts General Hospital for insightful discussions on human GWAS studies. We also thank Dr. Alex Banks and Marissa Cortopassi for their support in metabolic cage studies at the BIDMC metabolic core. Lastly, we thank Anthony R. P. Verkerke, Daisuke Kato, Tadashi Yamamuro, Martin Charles in the Kajimura lab for their technical help. This work was supported by the NIH (DP1DK126160) and the Howard Hughes Medical Institute to S. K. The work is also supported by DK081418 (P.P.), DK117655 (P.P.), 1F32GM136019–01A1 (B.M.), R00DK117066 (S.H.), and the Paul G. Allen Family Foundation 0034665 (S.H.).

## METHOD

### Animal study

All the animal experiments in this study were performed in compliance with protocols approved by the Institutional Animal Care and Use Committee (IACUC) at Beth Israel Deaconess Medical Center. The *Slc25a47* floxed (*Slc25a47*^flox/flox^) mouse was generated by *in vitro* fertilization of homozygous sperm (UC David) from Slc25a47^tm1a (EUCOMM)Hmgu^ targeting exons 5 and 6 of the *Slc25a47* gene in C57BL/6J background. A floxed LacZ -neomycin cassette on the Tm1a allele was removed using a FLP/Frt deletion by breeding *Slc25a47*^flox/flox^ with FLP deleter mice (Jackson Laboratory, Stock No. 009086). Then, *Slc25a47*^flox/flox^ mice were bred with Albumin Cre mice (Jackson Laboratory, Stock No. 003574) to generate liver-specific *Slc25a47* KO (*Slc25a47* KO^Liver^). Mice were kept under a 12 hr:12 hr light-dark cycle at ambient temperature (22-23 °C) and had free access to food and water. Mice were maintained on a regular chow diet or fed with a high-fat diet (60% fat, D12492, Research Diets) starting from 6 weeks old age for 6 weeks. All mice were fasted for 6 hours before sacrifice.

### Acute SLC25A47 deletion by AAV-Cre

To generate acute *Slc25a47* KO mice, we injected 7 weeks old *Slc25a47*^flox/flox^ mice with 1.5 x 10^11^ genome copies of AAV8-TBG-Cre (Addgene, 107787-AAV8) or AAV8-TBG-null (control, Addgene, 105536-AAV8) through tail vein injection.

### Human SNP analyses

Data were obtained from the Type 2 Diabetes Knowledge Portal (type2diabetesgenetics.org) and reconstructed. We used the SLC25A47 gene as the primary locus and expanded 5,000 bp proximal and distal to the total gene distance in order to identify regions of interest that may be outside of the coding sequence, *i.e*., promoters or enhancers.

### Transcriptional analysis

scATAC-seq and ChIP-seq data were obtained from GEO (GSE111586 and GSE90533, respectively) and visualized using IGV (Integrative Genomics Viewer). For the analysis of *SLC25A47* gene expression in human tissues and single cells of the human liver, the data was obtained from Human Protein Atlas (https://www.proteinatlas.org/ENSG00000140107-SLC25A47/tissue and https://www.proteinatlas.org/ENSG00000140107-SLC25A47/single+cell+type/liver, respectively). The data for mouse *Slc25a47* expression in tissues was obtained from GTEx portal (https://www.gtexportal.org/home/gene/SLC25A47). From these data, RNA expression at the tissue and single cell level was reconstructed.

### ^13^C-glucose and ^13^C-lactate infusion study *in vivo*

Jugular vein catheters (Instech Labs) were implanted in the right jugular vein of 10-week-old mice (n=6 per group) under aseptic conditions. The catheter was connected to a vascular access button (Instech Labs) into which the tracer was infused. After one week of the recovery period, mice were fasted for 6 h, and then infused for 2.5 h with U-^13^C-glucose (0.2 M, CLM-1396), U-^13^C-sodium lactate (0.49 M, CLM-1579), 0.2 M ^13^C-alanine (0.2 M, CLM-2184-H), and U-^13^C-glycerol (0.1M, CLM15101), respectively, at 2-3 days interval. The infusion rate was 0.1 μL·g^−1^·min^−1^, and mice moved freely in a cage during the intravenous infusions. Blood (~ 10 μL) was collected from the tail into microvettes with coagulating activator (Starstedt Inc, 16.440.100). Blood samples were kept on ice, and serum was separated by centrifugation at 3,000 g for 10 min at 4 °C. 4 μL of serum was added to 60 μL of ice-cold extraction solvent (methanol: acetonitrile: water at 40:40:20), vortexed vigorously and incubated on ice for at least 5 min. The samples were centrifuged at 3,000 rpm for 10 min at 4 °C, and the supernatant was transferred to LC-MS tubes for analyses.

### Serum metabolite analysis by LC/MS

Chromatographic separation was achieved using XBridge BEH Amide XP Column (2.5 μm, 2.1 mm × 150 mm) with guard column (2.5 μm, 2.1 mm X 5 mm) (Waters, Milford, MA). Mobile phase A was water: acetonitrile 95:5, and mobile phase B was water: acetonitrile 20:80, both phases containing 10 mM ammonium acetate and 10 mM ammonium hydroxide. The linear elution gradient was: 0 ~ 3 min, 100% B; 3.2 ~ 6.2 min, 90% B; 6.5. ~ 10.5 min, 80% B; 10.7 ~ 13.5 min, 70% B; 13.7 ~ 16 min, 45% B; and 16.5 ~ 22 min, 100% B, with flow rate of 0.3 mL/ min. The autosampler was at 4°C. The injection volume was 5 μL. Needle wash was applied between samples using methanol: acetonitrile: water at 40: 40: 20. The mass spectrometer used was Q Exactive HF (Thermo Fisher Scientific, San Jose, CA), and scanned from 70 to 1000 *m/z* with switching polarity. The resolution was 120,000. Metabolites were identified based on accurate mass and retention time using an in-house library, and the software used was EI-Maven (Elucidata, Cambridge, MA). ^13^C-Natural abundance correction was performed in R using package AccuCor (50).

### Calculation of direct contribution fraction of gluconeogenic substrates to glucose

The calculation follows the method as prior reported (51, 52). Briefly, for a metabolite with carbon number *C*, the labeled isotopologue is noted as [*M* + *i*], and its fraction is noted as *L*_[*M + i*]_, with *i* being the number of ^13^C atoms in the isotopologue. The overall ^13^C labeling *L_metabolite_* of the metabolite is calculated as the weighted average of atomized labeling of all isotopologues, or mathematically

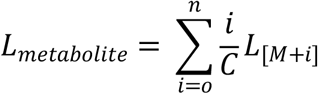

The normalized labeling *L_metabolite←tracer_* is defined as the labeling of a metabolite normalized by the labeling of the infused tracer, as

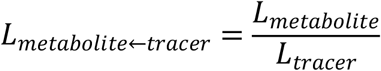

As such, the direct contribution of gluconeogenic substrates to glucose production is algebraically calculated by solving the matrix equation

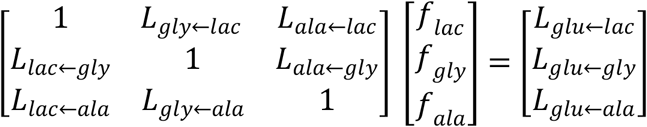

Specifically, let ***M*** be the matrix and ***f*** the vector on the left side, and ***L*** the vector on the right side. The operation seeks to

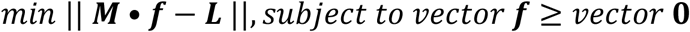

The equation is solved using the R package *limSolve* (53). The error was estimated using Monte Carlo simulation by running the matrix equation 100 times, each time using randomly sampled *L_metabolite←tracer_* values drawn from a normal distribution based on the mean and standard error of entries in ***M*** and ***f***. The calculated ***f***’s were pooled to calculate the error.

This scheme was extended to calculate the mutual interconversions among the metabolites.

### Calculation of normalized ^13^C enrichment of metabolites

The peak intensity of each measured isotope was corrected by natural abundance. To calculate the fraction of ^13^C labeled carbon atoms of glucose, pyruvate, lactate, glutamine, and alanine derived from ^13^C glucose and ^13^C lactate, percent ^13^C enrichment (%) was first calculated from the data corrected by natural abundance and then normalized based on the serum tracer enrichment.

### Mitochondrial isolation

Mitochondria were isolated from the fresh liver of 7-week-old mice following the modified protocol of Frezza et al. (54). Buffer for cell and mouse liver mitochondria isolation (IBc) (200mM sucrose, 10mM Tris/MOPS, 1mM EGTA, pH 7.4) was prepared fresh before each experiment. Animals were fasted for 6 hours, starting between 8 and 11 AM. To remove blood, we perfused the liver with 10 mL of ice-cold IBc by injecting slowly into the heart. After perfusion, the large left lobe of the liver was isolated and washed briefly with ice-cold IBc. The liver was transferred to a glass dounce homogenizer pre-filled with 8 mL of IBc buffer. All the steps were performed on ice or in a 4 °C cold room. The liver was homogenized using a Teflon pestle (Fisher Scientific, FSR3000) by 5 strokes at 1,500 rpm. The homogenate was then transferred into a pre-chilled 15 mL conical tube and spun at 600 g for 10 min at 4 °C. The supernatant, containing mitochondria, was transferred to a 15 mL tube and centrifuged at 600 g for 10 min at 4 °C, and then the supernatant was re-centrifuged 7,000 g at 4 °C in a fixed rotor centrifuge for 10 min for complete separation. The supernatant was removed, and the pellet containing mitochondria was washed with 1 mL of ice-cold IBc, and centrifuged at 7,000 g for 10 min at 4 °C. After discarding the supernatant, the pellet was resuspended with 1 mL of IBc and spun at 8,000g for 10 min at 4 °C. To remove IBc buffer, the mitochondrial pellet was gently washed twice by gentle resuspension in modified KPBS buffer (136 mM KCL, 10 mM KH2PO4, 10 mM HEPES, pH 7.25) and spun at 10,000 g for 60 s at 4 °C and repeated washing step for a total of two washes. After the second wash, we transferred 1 mL of the resuspended mitochondria into a new Eppendorf tube and then spun down the mitochondria at 10,000 g for 60 s at 4 °C. After centrifugation, we removed 750 uL of supernatant and then gently resuspended the pellet in the around 200 uL of remaining KPBS buffer to make a homogenous mixture. 50 uL of mitochondria suspension was used for mitochondrial metabolomics studies and tracing study with ^13^C pyruvate, and 10 uL of mitochondria was used for protein quantifications by BCA Assays and Western blots.

### ^13^C-tracers in the liver mitochondria

Fifty microliters of isolated mitochondrial suspension that was prepared as above was added into 450 uL of KPBS containing 2 mM U-^13^C pyruvate (Cambridge Isotopes, CLM-2440-0.1) and then incubated on ice for 5 min to inactivate enzymes for pyruvate uptake analysis or at 37 °C with gentle rotation at 300 rpm for 15 min for pyruvate flux analysis in mitochondria. After incubation, samples were immediately centrifuged at 10,000 g x 30 s, and then gently washed 3 times by adding 1 mL of ice-cold KPBS. After the final wash, the supernatant was removed and 1 mL of ice-cold LC/MS 80% methanol was added. After pipetting up and down, the sample was vortexed vigorously for 15 s before placing it on dry ice to let metabolites extract for 5 minutes and then kept in −80 °C until further analysis. To completely extract metabolites from the mitochondria, the sample was homogenized using TissueLyser II (Qiagen, 85300) for 5 min at 30 Hz, followed by centrifugation at 20,000 g for 15 min at 4 °C. The supernatant was saved and kept on dry ice, and the pellet was resuspended with 500 uL of 80% LC/MS-grade methanol, vortexed vigorously, and allowed to extract on ice. The samples were then centrifuged at 20,000 g for 10 min at 4 °C. The supernatants were pooled in a 2 mL Eppendorf tube. The pooled supernatants were then spun at 20,000g for 10 min at 2 °C, and the supernatants were transferred to a new tube on dry ice. The extraction was vacuum dried using a vacuum concentrator (Eppendorf, concentrator Plus 5305) and kept at −80 °C until metabolomics analyses.

### Metabolites analyses in mitochondria

Dried samples prepared as above were solubilized in 50 uL of LC/MS water and vortexed vigorously. The samples were kept on ice and centrifuged at 17,000 g for 5 min at 4 °C to pellet any debris. Metabolite analysis was conducted at the BIDMC metabolomics core. The data were normalized by protein concentration.

### RNA-Sequencing

Extracted RNA (400ng) from control and *Slc25A47* KO^Liver^ liver samples were treated with NEBNext rRNA Depletion Kit v2 (E7400X) to remove ribosomal RNA. The first stand was synthesized using Thermo Scientific Random Hexamer Primer (SO142) and Maxima Reverse Transcriptase (EP0742). The second strand was synthesized using NEBNext mRNA Second Strand Synthesis Module (E6111L). cDNA was analyzed with Qubit and Agilent BioAnalyzer, and subsequently amplified for 12 cycles using Nextera XT DNA Library Preparation Kit (Illumina FC-131). Generated libraries were analyzed with Qubit and Agilent Bioanalyzer, pooled at a final concentration of 1.35 pM, and sequenced on a NextSeq 500. Sequenced reads were demultiplexed and trimmed for adapters using bcl2fastq (v2.20.0). Secondary adapter trimming, NextSeq/Poly(G) tail trimming, and read filtering was performed using fastp (v0.23.2) (55); low quality reads and reads shorter than 24nt after trimming were removed from the read pool. Salmon (v1.6.0) (56) was used to simultaneously map and quantify reads to transcripts in the GENCODE M26 genome annotation of GRCm39/mm39 mouse assembly. Salmon was run using full selective alignment, with sequence-specific and fragment GC-bias correction turned on (seqBias and gcBias options, respectively). Transcript abundances were collated and summarized to gene abundances using the tximport package for R (57). Normalization and differential expression analysis were performed using edgeR (58, 59). For differential gene expression analysis, genes were considered significant if they passed a fold change (FC) cutoff of log2FC < 1 and a false discovery rate (FDR) cutoff of FDR < 0.05.

### Glucose, insulin, pyruvate, and glycerol tolerance tests

For glucose and insulin tolerance tests, mice were fasted for 6 h from 9:00 to 15:00 prior to glucose (2 g/kg BW) or insulin (0.4 U/kg BW) injection intraperitoneally. For insulin measurement during the glucose tolerance test, blood was collected 15 min after the glucose administration. For pyruvate and glycerol tolerance tests, we fasted mice for 16 h and injected pyruvate (2 g/kg BW) or glycerol (2 g/kg BW) intraperitoneally. Blood glucose levels were measured at the indicated time points using blood glucose test strips (Freestyle Lite).

### Metabolic analysis

The metabolic variables, including oxygen consumption rate (VO_2_), carbon dioxide release rate (VCO_2_), energy expenditure (EE), food intake, and locomotor activity (beam break counts), were measured using the Promethion Metabolic Cage System (Sable Systems) at 23 °C at 1 week of a high-fat diet, and at 30 °C at 3 weeks of a high-fat diet. Body Composition (lean mass and fat mass) was measured using Analyzer EchoMRI (Echo Medical Systems). As for examining metabolic changes after CL-316,243 injection, mice on a regular chow diet were housed at 30 °C for one week prior to CL injection to acclimate to thermoneutrality. The metabolic variables were measured using the Promethion Metabolic Cage System at 30 °C before and after CL-316,243 (0.5mg/kg weight *i.p*.) injection.

### Serum analysis

Blood was coagulated at room temperature for 1 h, and then serum was separated by centrifugation at 2,000 g for 20 min at 4 °C and kept at −80 °C until further analysis. Total cholesterol (TC, Stanbio), triglyceride (TG, Thermos Fisher), insulin (Millipore Sigma), FGF21 (R&D system), AST and ALT (Abcam), and albumin (Stanbio) were analyzed in the serum according to the manufacturer’s instruction.

### Histology

Liver samples were fixed in 4% paraformaldehyde (PFA) at 4 °C overnight, followed by dehydration in 70% ethanol. After dehydration, the samples were processed for paraffin embedding and cross-sectioned at 5 μm thickness. Sections were subjected to hematoxylin and eosin (H&E) staining or picro-sirius red staining according to the standard protocol. Images were acquired using the Zeiss AxioImager M1 (Carl Zeiss).

### Glycogen measurement

Glycogen content was determined using a commercially available kit (Abcam). Briefly, 10 mg of liver tissue was homogenized in cold distilled water using two stainless steel beads (TissueLyser II for 1 min at 40 Hz). The samples were then boiled at 95 °C for 10 mins to inactivate enzymes, and insoluble material was removed after centrifugation at 18,000 g at 4 °C for 10 mins. The supernatant was quantified by colorimetric assays, and glucose level was subtracted from the glycogen levels.

### Oxygen consumption rate

To isolate mitochondria, liver samples were homogenized in 3 mL of MSHE buffer (70 mM sucrose, 210 mM mannitol, 5 mM HEPES, 1 mM EGTA, pH 7.4) containing 0.5% free fatty acid free BSA on ice using a Teflon pestle (Fisher Scientific, FSR3000) by 5 strokes at 1,500 rpm. The homogenate was transferred into a cold 15 ml and centrifuged at 600 g for 10 min at 4 °C. The supernatant was then transferred into multiple cold 1.5 mL EP tubes and spun at 1,100 g for 10 min at 4 °C. The pellet was resuspended in 500 μL of MSHE and then centrifuged at 8,000 g for 10 min at 4 °C. To remove BSA for protein quantification, the pelleted mitochondria were washed once and resuspended in MSHE without BSA. Oxygen consumption rate in liver mitochondria was measured using the Oroboros Oxygraph 2k (NextGen-O2k, 10101-01). Briefly, 80 ug of mitochondria was added into the calibrated Oroboros chamber. To obtain state 4 respiration linked to CI, 5 mM pyruvate, 2 mM malate, and 10 mM glutamate were injected. State 3 respiration was measured via the addition of 4 mM ADP. Injection of 10 mM succinate permitted the measurement of CI and CII-driven respiration.

### qRT-PCR

Total RNA was isolated from the liver by using TRIzol Reagent (Invitrogen) and further purified using Zymo direct-zol RNA prep (Zymo research) according to the manufacturer’s protocol. Genomic DNA was removed using DNAse during RNA purification, and cDNA was synthesized using iScript^™^ Reverse Transcription Supermix (Biorad) following the provided protocol. RT-PCR was performed on a QuantStudio™ 6 (Life technologies) using iTaq^™^ Universal SYBR^®^ Green Supermix (Biorad). Relative mRNA levels were calculated based on the 2^−ΔΔct^ method with normalization of the raw data to *18s*. Primer sequences are available in supplementary table 1.

### Statistical analyses

Statistical analyses were performed using GraphPad Prism 9.4.1 (GraphPad Software) and R studio. All data were represented as mean ± SEM unless otherwise specified. Unpaired Student’s t-test and Mann-Whitney test were used for two-group comparisons. Two-way repeated-measures ANOVA followed by Fisher’s LSD test was applied to determine the statistical differences in body weight change, whole-body energy expenditure results, glucose tolerance test, insulin tolerance test, and pyruvate tolerance test between genotypes. Testing the direct association between two variables was performed using simple linear regression. The statistical parameters and mice numbers used per experiment are clarified in the figure legends. *p* < 0.05 was considered significant in all the experiments.

---

**Supplementary Figure S1.**
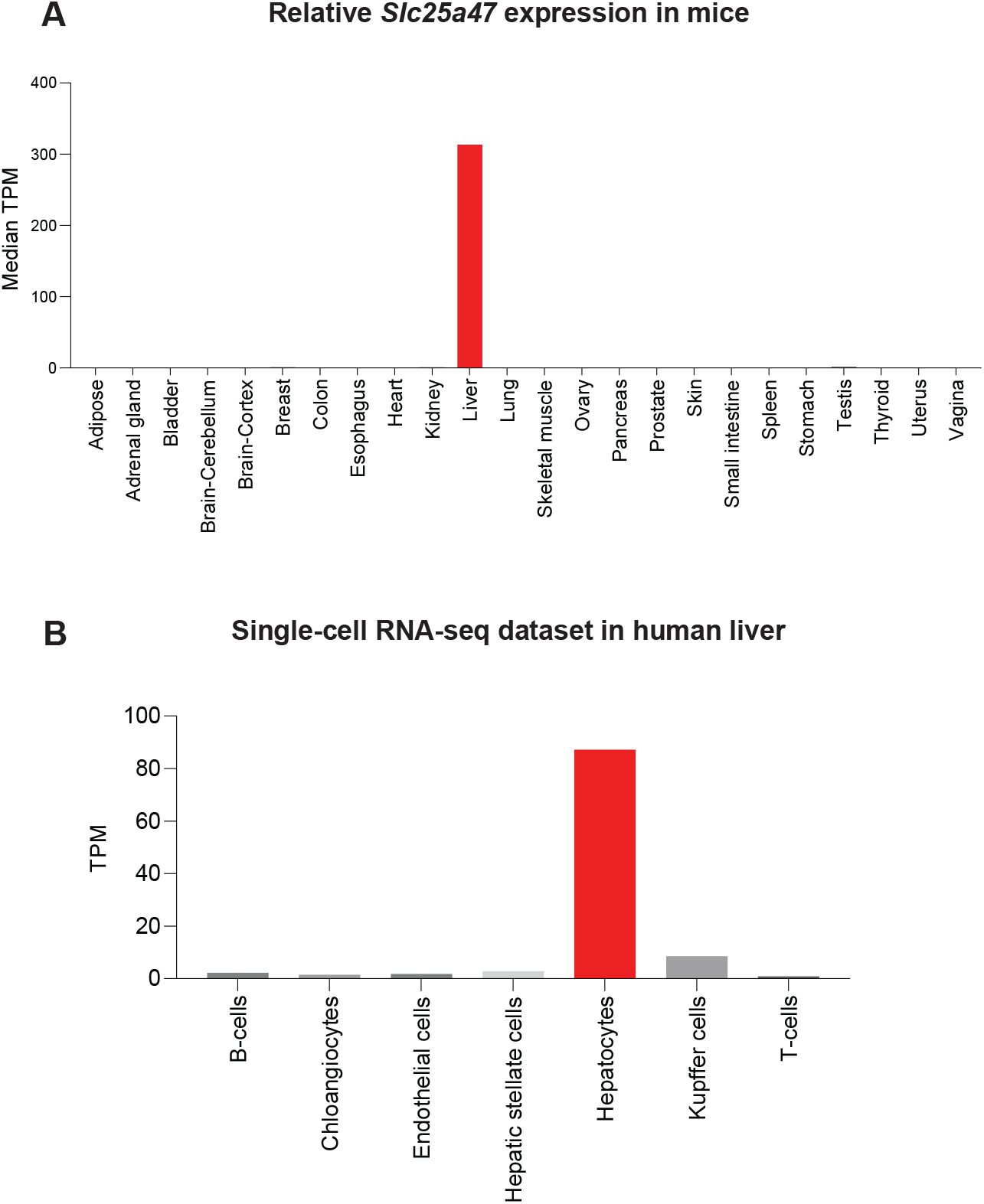
SLC25A47 is a liver-specific mitochondrial carrier in mice. **A.** Relative mRNA levels (TPM) in indicated tissues of mice. The data obtained from GTEx portal (https://www.gtexportal.org/home/gene/SLC25A47) were analyzed. **B.** Relative mRNA levels (TPM) in indicated cell types in the human liver. The data obtained from https://www.proteinatlas.org/ENSG00000140107-SLC25A47/single+cell+type/liver were analyzed.

**Supplementary Figure S2.**
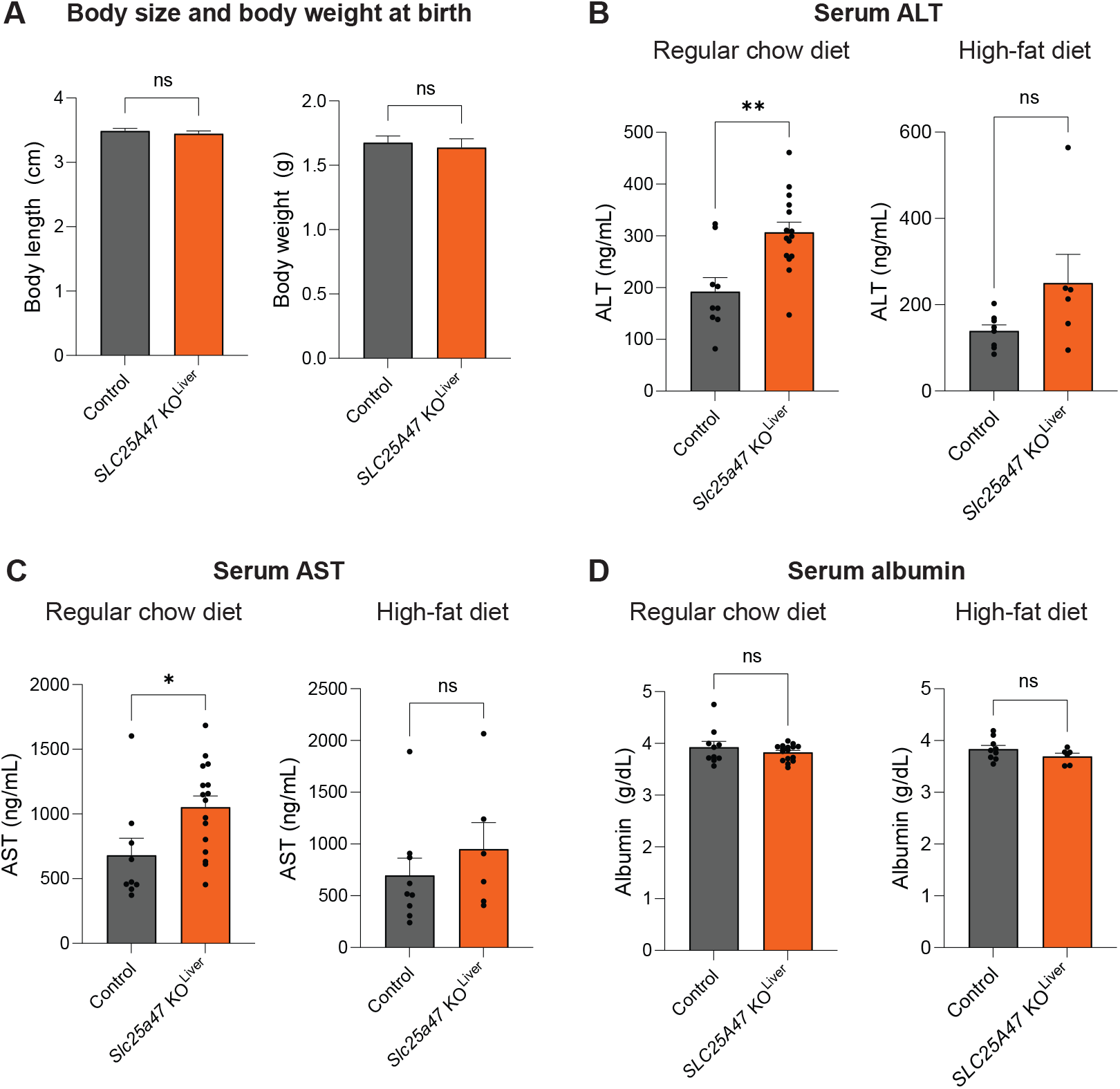
Characterization of *Slc25a47* KO ^Liver^ mice. **A.** Body length and body weight in *Slc25a47* KO ^Liver^ mice and littermate control mice in newborn. *n* = 10 for *Slc25a47* KO ^Liver^ mice, *n* = 18 for controls. Data are mean ± SEM.; ns, not significant, by twotailed unpaired Student’s *t*-test. **B.** Serum ALT levels of *Slc25a47* KO ^Liver^ mice and littermate control mice on a regular chow diet (left) and high-fat diet (right). *n* = 16 for *Slc25a47* KO ^Liver^ mice, *n* = 10 for controls (regular chow diet), *n* = 6 for *Slc25a47* KO ^Liver^ mice, *n* = 9 for controls (high-fat diet). Data are mean ± SEM.; * *p* < 0.05, ** *p* < 0.01, ns, not significant, by two-tailed unpaired Student’s *t*-test. **C.** Serum AST levels of mice in (B). **D.** Serum albumin levels of mice in (B).

**Supplementary Figure S3.**
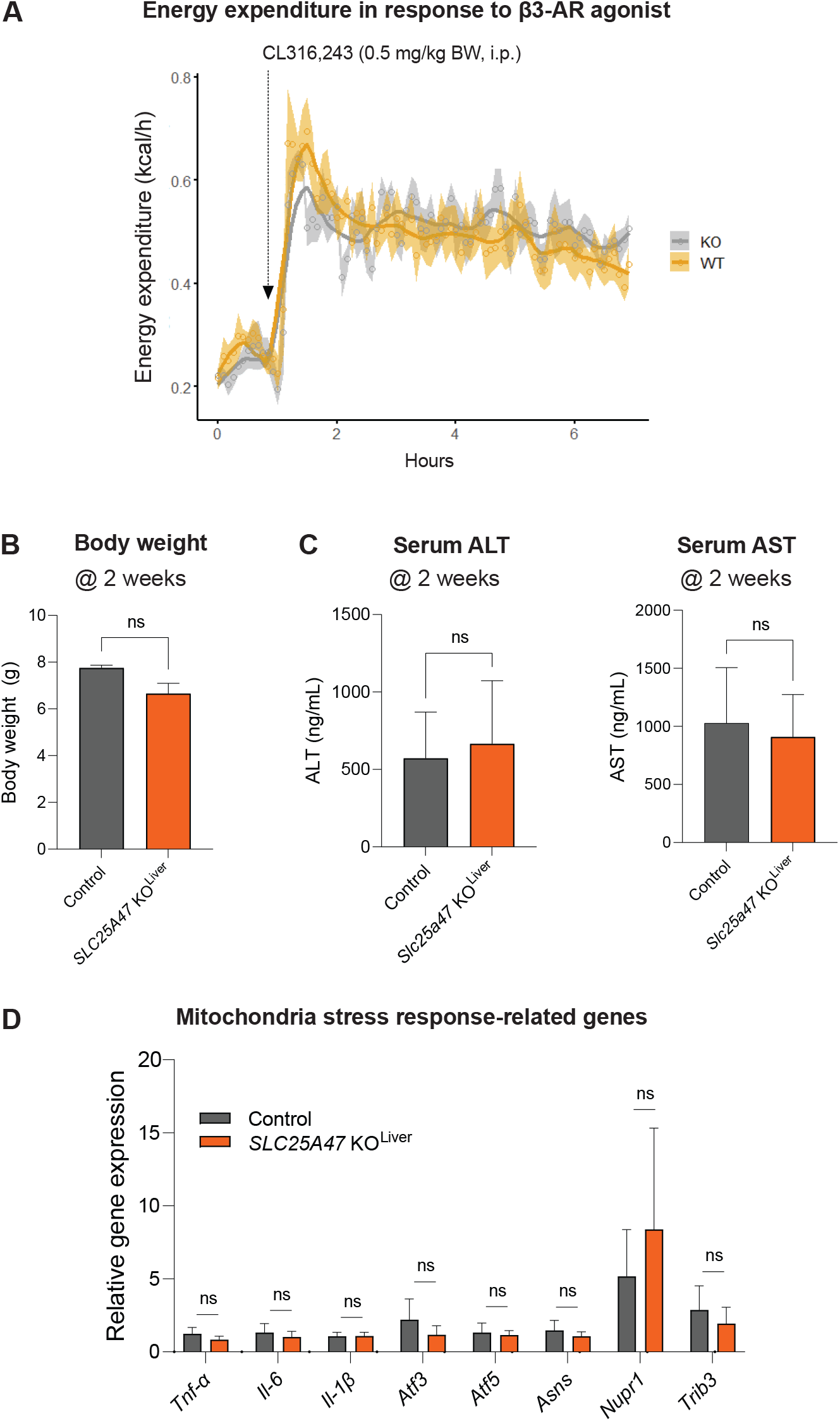
Brown adipose tissue thermogenesis and liver function of *Slc25a47* KO ^Liver^ mice. **A.** Energy expenditure (kcal/h) of *Slc25a47* KO ^Liver^ mice and littermate controls on a regular chow diet. Mice kept at 30°C received *i.p*. injection of CL316,243 at 0.5 mg kg^-1^ at the indicated time point (arrow). *n* = 8 for both groups, biologically independent mice. Data are mean ± SEM.; *P*-value was determined by two-way repeated-measures ANOVA. **B.** Body weight in *Slc25a47* KO ^Liver^ mice and littermate control mice at 2 weeks age. *n* = 6 for *Slc25a47* KO ^Liver^ mice, *n* = 4 for controls. Data are mean ± SEM.; ns, not significant, by two-tailed unpaired Student’s *t*-test. **C.** Serum ALT (left) and AST (right) levels of *Slc25a47* KO ^Liver^ mice and littermate control mice at 2 weeks of age in (B). Data are mean ± SEM.; ns, not significant, by two-tailed unpaired Student’s *t*-test. **D.** Expression levels of genes related to inflammation and mitochondrial stress-response in *Slc25a47* KO ^Liver^ mice and littermate control mice at 2 weeks age in (B). *n* = 6 for *Slc25a47* KO ^Liver^ mice, *n* = 4 for controls. Data are mean ± SEM.; ns, not significant, by two-tailed unpaired Student’s *t*-test.

**Supplementary Figure S4.**
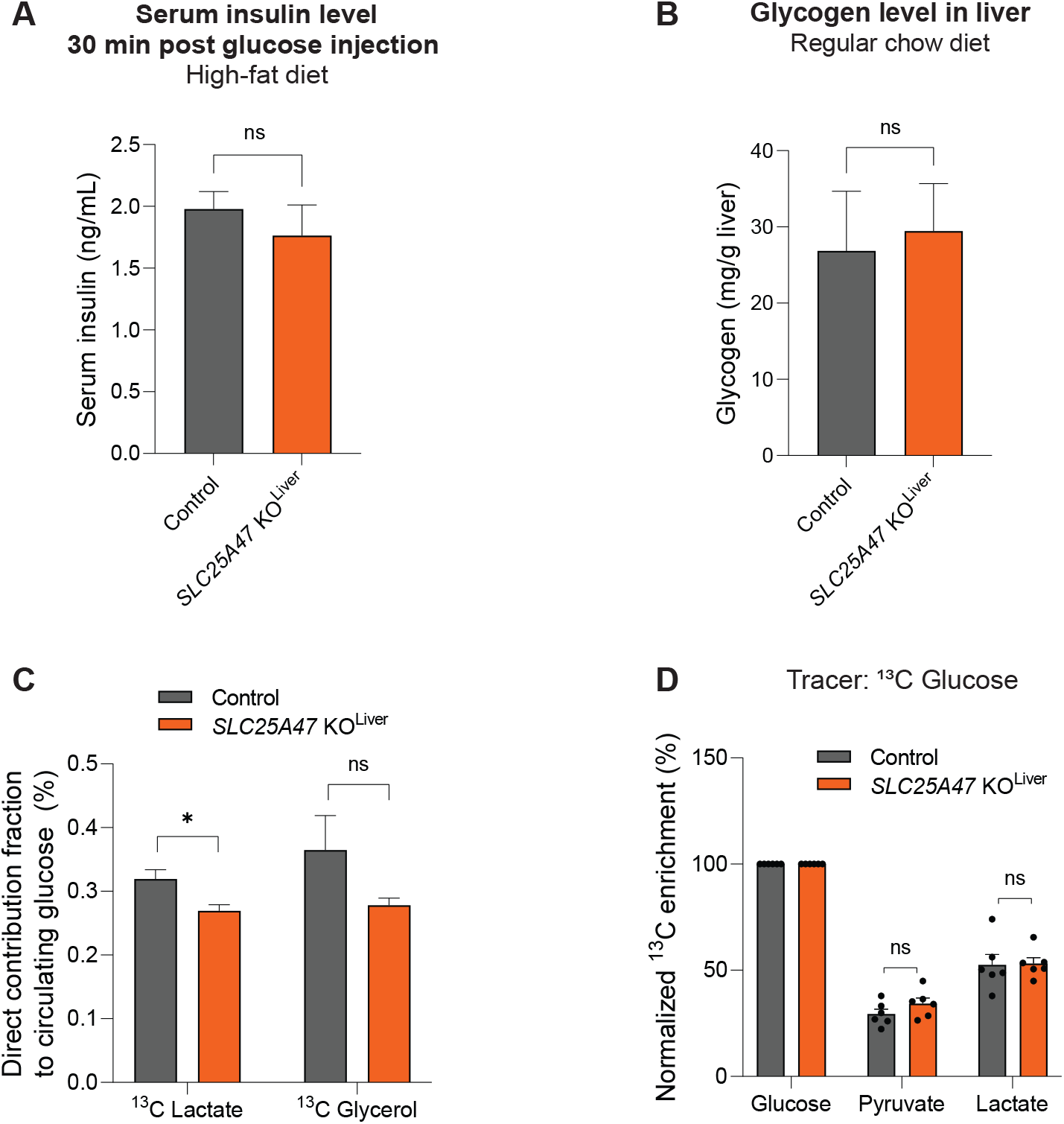
Glucose metabolism of *Slc25a47* KO ^Liver^ mice. **A.** Glucose-stimulated serum insulin levels at 30 minutes after glucose injection in *Slc25a47* KO ^Liver^ mice and littermate control mice on a high-fat diet (HFD). *n* = 10 for *Slc25a47* KO ^Liver^ mice, *n* = 5 for controls. Data are mean ± SEM.; ns, not significant, by two-tailed unpaired Student’s *t*-test. **B.** Glycogen levels in the liver of *Slc25a47* KO ^Liver^ mice and littermate control mice at 12 weeks of age on a regular chow diet (RD). *n* = 16 for *Slc25a47* KO ^Liver^ mice, *n* = 9 for controls. Data are mean ± SEM.; ns, not significant, by two-tailed unpaired Student’s *t*-test. **C.** Direct contribution of ^13^C-labeled glucose and glycerol to circulating glucose. *n* = 6 for *Slc25a47* KO ^Liver^ mice, *n* = 6 for control mice, biologically independent mice. Data are mean ± SEM.; * *p* < 0.05, ns, not significant, by unpaired Student’s *t*-test. **D.** ^13^C-enrichment (%) of labeled glucose to circulating levels of indicated metabolites in (C). ns, not significant, by unpaired Student’s *t*-test.

**Supplementary Figure S5.**
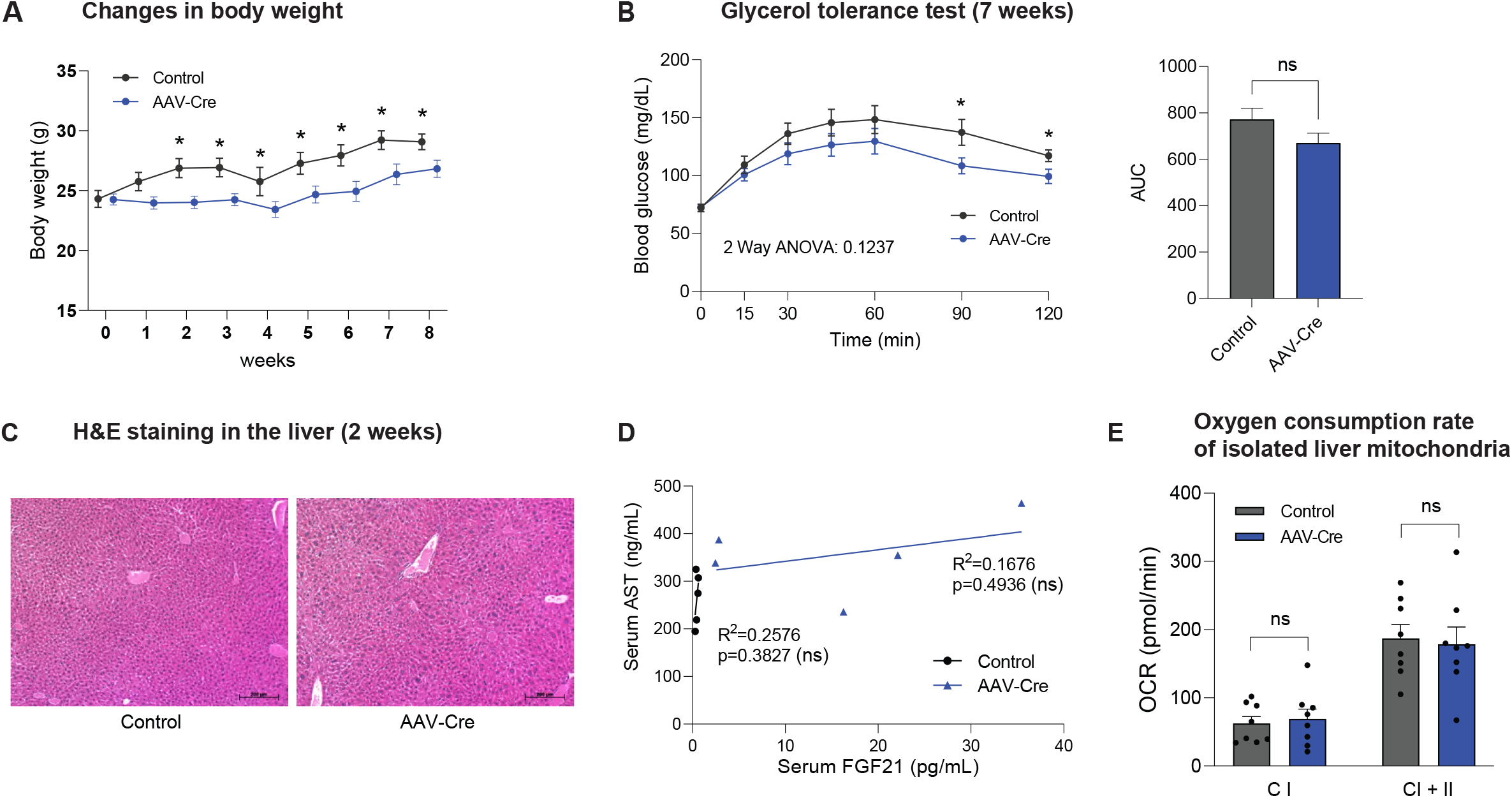
Metabolic changes in response to acute SLC25A47 depletion in adult mice. **A.** Changes in body weight of *Slc25a47*^flox/flox^ mice that received AAV-Cre or AAV-null (control) at indicated time points. Mice were on a regular chow diet. *n* = 10 for controls, *n* = 13 for AAV-Cre, biologically independent mice. Data are mean ± SEM.; *P*-value was determined by unpaired Student’s *t*-test. **B.** Glycerol tolerance test in *Slc25a47*^flox/flox^ mice at 7 weeks after receiving AAV-Cre or AAV-null (control). After 16 hours of fasting, mice received *i.p*. injection of glycerol at 2 g kg^-1^ body-weight. *n* = 11 for controls, *n* = 14 for AAV-Cre, biologically independent mice. Data are mean ± SEM.; *P*-value was determined by two-way repeated-measures ANOVA followed by Fisher’s LSD test. Right: AUC of the data was calculated by Graphpad software. ns, not significant, by two-tailed unpaired Student’s *t*-test. **C.** Representative image of hematoxylin and eosin (H&E) staining in the liver of *Slc25a47*^flox/flox^ mice at 2 weeks after receiving AAV-Cre or AAV-null (control). Scale = 200 μm. **D.** Correlation between serum FGF21 levels and serum AST levels in *Slc25a47*^flox/flox^ mice at 2 weeks after receiving AAV-Cre or AAV-null (control). *n* = 5 for controls, *n* = 5 for AAV-Cre, biologically independent mice. ns, not significant **E.** Oxygen consumption rate (OCR) of isolated mitochondria from the liver of *Slc25a47*^flox/flox^ mice that received AAV-Cre or AAV-null (control). Mitochondrial Complex I (CI) activity was measured by adding glutamate (10 mM), pyruvate (5 mM), and malate (2 mM). Succinate (10 mM) was then added to measure the respiration of complexes I and II. *n* = 8 for controls, *n* = 8 for AAV-Cre, biologically independent samples. Data are mean ± SEM.; ns, not significant, by two-tailed unpaired Student’s *t*-test.

**Supplementary Figure S6.**
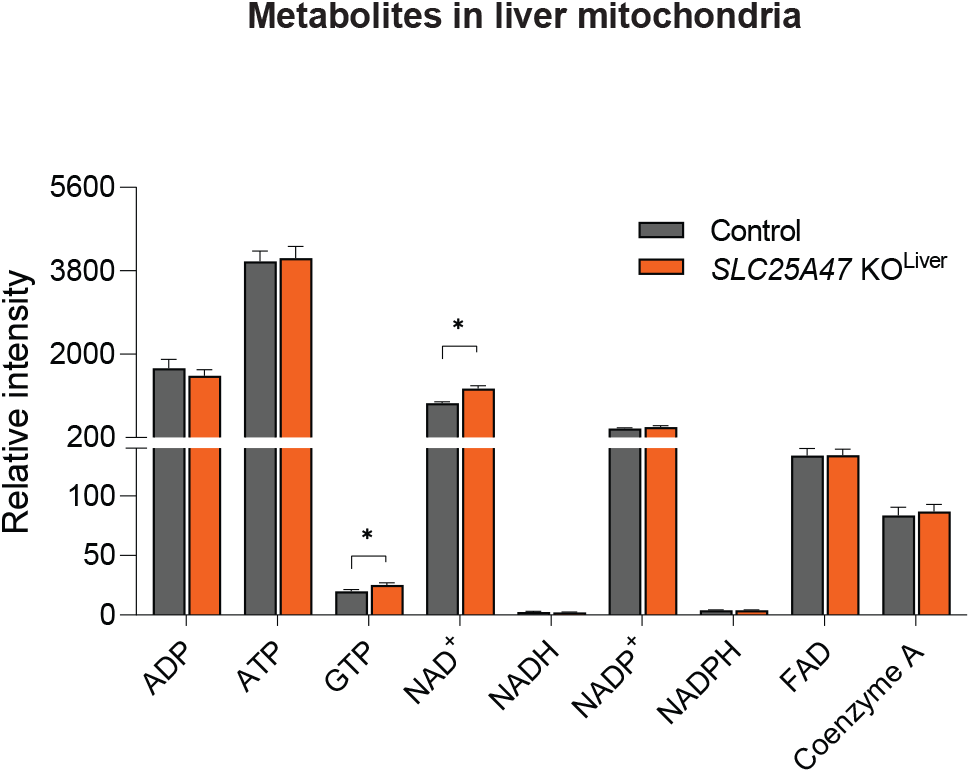
Mitochondrial metabolomics of the liver. Relative levels of indicated metabolites in the liver mitochondria from *Slc25a47* KO ^Liver^ mice and littermate controls in Fig. 5C. Mice at 7 weeks of age fasted for 6 hours. *n* = 14 for *Slc25a47* KO ^Liver^ mice, *n* = 14 for the littermate control mice, biologically independent mice. Data were normalized to mitochondrial protein levels and shown as mean ± SEM.; *P*-value was determined by two-tailed unpaired Student’s *t*-test.

**Supplemental Table 1.**
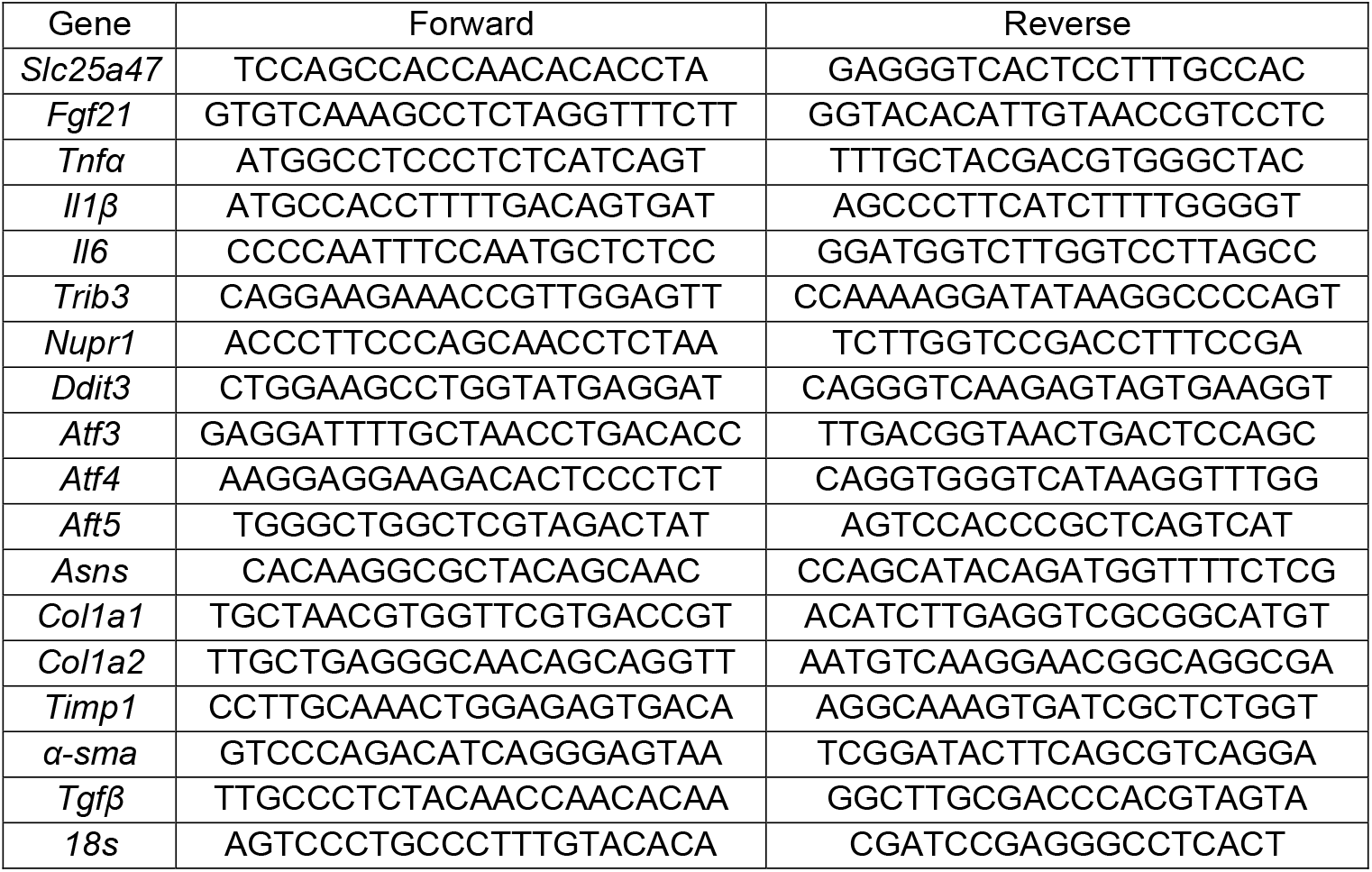
List of mouse primer sequences for qPCR.

## Notes

### Competing Interest Statement

The authors have declared no competing interest.

